# CombDNF: Disease-specific drug combination predictions with network-based features on clinically validated data

**DOI:** 10.1101/2025.02.18.637825

**Authors:** Pauline Hiort, Bernhard Y. Renard, Katharina Baum

**Affiliations:** Hasso Plattner Institute, Digital Engineering Faculty, University of Potsdam, Potsdam, Germany; Institute of Computer Science, Department of Mathematics and Computer Science, Free University Berlin, Berlin, Germany; Hasso Plattner Institute for Digital Health at Mount Sinai and Windreich Dept. of Artificial Intelligence & Human Health, Icahn School of Medicine at Mount Sinai, New York, USA

## Abstract

Drug combinations are increasingly applied to treat a wide range of complex diseases. Drug action and thus also drug combination effects can differ between diseases, e.g., due to molecular differences. Therefore, disease-specific predictions are required for treatments. A plethora of methods based on cell-line screening data in cancer has been proposed. However, their extendability to other diseases is limited, as is their applicability in the clinical context due to the in-vivo-in-vitro gap. In contrast, only few approaches rely on clinically validated data.

Here, we propose CombDNF, a novel machine-learning based method for disease-specific drug combination prediction on clinically validated data. CombDNF is trained and predicts both on clinically approved (effective) and clinically reported adverse drug combinations from a broad collection of data sources. It can cope with the highly imbalanced label distribution in drug combination data. Further, CombDNF leverages network-derived features based on drug target and disease gene relationships. To incorporate uncertainty of the network topology it relies on edge weights in the underlying network.

We systematically evaluate CombDNF against available state-of-the-art methods in four diseases with different underlying mechanisms and ground truth data characteristics. CombDNF outperforms all state-of-the-art methods in all four diseases by at least 84% in the AUPR. This translates, for example, to an enrichment of effective drug combinations in the top ten hypertension-specific predictions of four, compared to one for the best competing method. In addition, network edge weighting by interaction confidence scores indeed yields improved predictions. Further, we find evidence for biological plausibility of our top ranked drug combinations.

We make the training and evaluation pipeline for CombDNF available ready-to-use at https://github.com/DILiS-lab/CombDNF.

## Introduction

Combinations of drugs are increasingly applied to treat complex diseases, such as cancer, cardiovascular disorders, and infectious diseases (Das et al., 2019). Combining drugs increases the number of targets in diseases which can lead to greater therapeutic effects, overcome drug resistance, and reduce toxicity compared to single drugs (Güvenç Paltun et al., 2021; Madani Tonekaboni et al., 2018). On the other hand, administering drug combinations can lead to drug-drug interactions and, thereby, to unexpected new and adverse effects. Analyzing all possible combinations of two drugs vastly increases the search space for suitable treatments. Computational methods have been proposed to reduce this space to a few candidate combinations, e.g., reviewed in (Güvenç Paltun et al., 2021; Madani Tonekaboni et al., 2018; Sheng et al., 2018).

Further, it is well established that drug action is highly disease-specific, and distinguishing responding from non-responding cohorts has been a key task in drug-related research. One underlying reason is the disease-specificity of protein expression and protein activity as mediators of drug action (Alvarez et al., 2016). Some sources, such as PharmGKB (Whirl-Carrillo et al., 2021), consider differences in mutation profiles and relate them to drug effect and side effects. Analogously, the effect of drug combinations can differ when applied to patients with different diseases. In cancer treatment, for example, disease-selectivity is crucial to predict drug combinations that target diseased cells while not affecting healthy cells (Kong et al., 2022). Therefore, it is vital to resolve drug combination predictions on a disease-specific level (Sheng et al., 2018; Madani Tonekaboni et al., 2018).

In the last decades, large-scale, high-throughput *in vitro* drug and drug combination screenings were performed to gain further (disease-specific) insights. These were mainly conducted using different cancer cell lines or infectious disease models (Liu et al., 2023; Sheng et al., 2018; Wu et al., 2022). Based on this screening data as ground truth, a plethora of computational methods predicting drug interactions in a cell-line specific but also unspecific manner have been proposed, e.g., reviewed in Wu et al. (2022); Kong et al. (2022); Liu et al. (2023); Chen et al. (2025). Predictions relying on screening results in cell lines and infectious disease models have two key shortcomings. First, especially large and systematic screenings are limited to diseases with appropriate simple cell line models, such as cancer. Thus, for drug combinations, e.g., in systemic diseases that lack cell line models, commonly no larger-scale screening data is available (Horvath et al., 2016; Wu et al., 2022). Second, there is the *in-vivo-in-vitro* gap where cell-line-based models can lack representativeness for the entire *in vivo* system (Horvath et al., 2016; Jaroch et al., 2018; Madani Tonekaboni et al., 2018). For these reasons, it is difficult to derive clinically relevant predictions from screening data (Madani Tonekaboni et al., 2018).

A preferable approach relies on clinically validated drug combination data. These are drug combinations that have officially been approved for patient treatment by governmental institutions, such as the U.S. Food and Drug Administration (FDA) (e.g., from DCDB (Liu et al., 2014) or CDCDB (Shtar et al., 2022)), or that have clinically reported adverse effects (e.g., from DrugBank (Wishart et al., 2018) or TWOSIDES (Tatonetti et al., 2012)). Clinically validated data have been used with computational methods to predict or evaluate effective drug combinations, e.g., Iwata et al. (2015); Liu et al. (2019); Zhao et al. (2011); Jaeger et al. (2017), side effects of drug combinations, e.g., Zitnik et al. (2018), or effective and adverse drug combinations, e.g., Yu et al. (2022); Gu et al. (2023). Thereby, some methods have leveraged disease-specific information to characterize the drugs for their drug interaction predictions, e.g., Yoo et al. (2018). However, the majority of published prediction methods do not aim to resolve differences in drug combination effects between diseases (Wu et al., 2022).

Only few methods have been developed for disease-specific (effective) drug combination prediction on clinically validated data for non-communicable diseases so far (see Table 1 for a summary). We identified these seven disease-specific drug combination characterization and prediction methods by systematically searching the literature. In particular, we screened all publications cited in relevant reviews (Wu et al., 2022; Madani Tonekaboni et al., 2018; Güvenç Paltun et al., 2021), and all publications that cite (as of Oct 1st, 2024) either one of the two systematic databases capturing FDA-approved, clinically validated drug combinations (Liu et al. (2014) and Shtar et al. (2022)). We included all those methods that provided disease-specific predictions or scores for effective drug combinations (for non-communicable diseases) if they relied on clinically validated drug combination data. We excluded purely cell line-based prediction methods and drug combination side effect predictions as these have a different prediction objective.

**Table 1.** Overview of published methods for disease-specific prediction of effective drug combinations based on clinically validated data in non-communicable diseases. We list if the methods are supervised (sup.), which underlying model is used, what types of drug combination predictions are possible (and their ground truth source, if applicable), the molecular basis of the model, how disease-specificity is achieved, whether it is extendable to other diseases (ext.), whether the descriptions are reproducible (repr.), and whether code is available. GE: gene expression, CNV: copy number variations, SMILES: chemical structures in Simplified Molecular Input Line Entry System, LINCS: Library of Integrated Network-Based Cellular Signatures

| Method | sup. | (ML) model | predictions | molecular basis | disease-specificity | ext. | repr. | code |
| --- | --- | --- | --- | --- | --- | --- | --- | --- |
| Pang et al. (2014) | no | mixed integer linear programming | effective only | drug target - disease gene overlap | disease genes | ✓ | — | — |
| Cheng et al. (2019) | no | PPI network proximity measures | effective & adverse (FDA + others) | drug targets - disease gene distances in PPI | disease genes | ✓ | ✓ | ✓ |
| CPGD (Guo et al., 2021) | no | nonlinear structural network controllability | effective & side effects separately | drug similarity from gene-gene network, LINC, SMILES | personalized network from GE of normal vs. tumour samples, CNV | — | (✓) | (✓) |
| Federico et al. (2022) | no | genetic algorithm | effective only | drug similarity in gene-gene knowledge graph, LINC, SMILES, drug targets | GE, LINC | (✓) | (✓) | ✓ |
| MCDC (Chen et al., 2019) | yes | kernel regularized least squares for link prediction | effective (DCDB), adverse (random) | SMILES, drug target properties, phenotype-based disease similarity | link in drug-pair-disease network | ✓ | — | — |
| VGAETF (Shan et al., 2023) | yes | graph variational auto-encoder | effective (CDCDB), adverse (random) | drug-drug-disease-disease network | link in drug-disease network | ✓ | ✓ | ✓ |
| KG-CombPred (Ye et al., 2023) | yes | graph attention neural network on knowledge graph | effective (literature), adverse (random) | PPI, drug targets, protein-GO network | infectious diseases-specific knowledge graphs | — | (✓) | (✓) |

Overall, as the molecular basis of disease and drug action, most methods rely on molecular networks as well as drug targets. We find that methods conferring disease-specificity via disease genes are extendable to other diseases. We further observe that methods are either unsupervised or learn based on random sampling among non-effective drug combinations for the negative class. The latter has two consequences: First, randomly sampled, negative class training data may inadvertently contain effective drug combinations. Second and more indirectly, the task of drug combination prediction is highly imbalanced, with decisively less effective than adverse combinations (Wu et al., 2022). Thus, as negative samples are generated at an effective-to-adverse ratio of 1:1 for published methods, the real-world label distribution is not well represented. This can lead to an over-optimistic estimation of model performance and a lack of generalization ability of trained models.

Based on these observations, to improve upon the state-of-the-art, we develop a novel supervised disease-specific drug combinations classification method, CombDNF (predicting drug **Comb**inations with **D**isease-specificity and **N**etwork-based **F**eatures). CombDNF trains on and predicts effective together with adverse clinically validated drug combinations. Thereby, the two classes are directly learned to be distinguished, and the model is evaluated under a more realistic label distribution scenario. In turn, this enhances its applicability for inference under realistic conditions. CombDNF extends features from network-based integration of drug- and disease-based information that were proposed for distinguishing both classes in Cheng et al. (2019). We rely on this approach as it was used manifold in related tasks, e.g., for drug molecular structure prediction (Karimi et al., 2020). However, uncertainty in the molecular interaction network topology is known to hamper conclusions (Alanis-Lobato et al., 2016). Therefore, we design CombDNF to improve on the earlier approach by supporting weighted instead of only binary molecular interaction networks for feature computation. Consequently, CombDNF can leverage confidence scores of molecular interactions to represent uncertainty in the network topology in its predictions. CombDNF is broadly generalizable to a wide variety of different diseases as it relies on sets of disease genes to make its predictions disease-specific. Finally, we choose eXtreme gradient tree boosting (XGB) as the prediction model for CombDNF. XGB is considered current state-of-the-art for small tabular datasets and has been used for similar tasks (Shwartz-Ziv and Armon, 2022; Liu et al., 2019). For the setup of training, prediction, and evaluation in CombDNF, we provide a Python pipeline that can be used for ML-based learning with arbitrary features for benchmarking drug combination classification. Further, we collect and provide ground truth data from several different sources for disease-specific, clinically relevant supervised drug combination classification. The ground truth data consists of both effective (approved as well as experimentally validated) and clinically reported adverse drug combinations.

For an evaluation from different perspectives, we compare CombDNF with state-of-the-art methods for disease-specific drug combination prediction from clinically validated data in four diseases: neoplasms (cancer), hypertension, cardiovascular, and nervous system diseases. These are selected to represent different characteristics especially in terms of frequency of occurrence, affected organ system, and disease mechanism. Neoplasms are highly heterogeneous with abnormal cell growth affecting different organs and tissues. For them, intracellular functions tend to play a crucial role. Cardiovascular diseases, and hypertension as a subtype of them, are often systemic, i.e., they affect multiple organs. They can be caused by a variety of (sub-)microscopic or macroscopic dysfunctions that lead to a similar phenotype. Compared to cancer, these diseases are more frequent, as are nervous system diseases. Underlying reasons of the latter are heterogeneous, ranging from mutations of a single gene to systemic causes. Their unifying characteristics is their effect on (parts of) the system related to processing of internal or external signals.

Further, we assess design choices of CombDNF, such as the effect of confidence score-based network edge weights and the single-disease models vs. an all-disease model approach.

In addition, we explore the influence of ground truth data characteristics, especially the observed label imbalance in this task, on the prediction performance. Finally, we investigate the biological plausibility of the predictions of CombDNF.

## Methods

We develop a novel method, CombDNF, for disease-specific drug combination classification on clinically validated data with network-based features. CombDNF includes two parts: network-based feature generation, and drug combination classification with XGB. During training and evaluation, we focus on reproducibility and evaluation avoiding data leakage. Further, our approach and code can also be used to benchmark drug-drug classification methods with arbitrary features. An overview of CombDNF is shown in Figure 1. Additionally, we collect and provide disease-specific ground truth data from different databases and literature. The ground truth data consist of experimentally validated, clinically relevant effective drug combinations, and adverse combinations with clinically reported adverse drug-drug interactions. For nuanced evaluation from different perspectives, four different disease groups are investigated that represent different levels of action (systemic vs. cellular), frequency of occurrence in the population (very frequent to less frequent), and affected organ systems (nervous system vs. various vs. blood vessels and heart): neoplasms (cancers), cardiovascular diseases, hypertension as a specialization of cardiovascular diseases, and nervous system diseases.

**Fig. 1:**
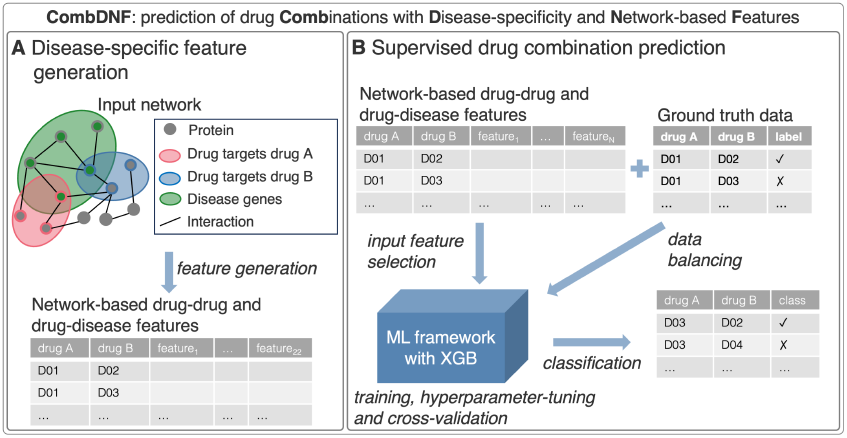
Overview of CombDNF for disease-specific drug combination classification with network-based features. (A) Generation of disease-specific features using a PPI network, drug targets, and disease genes. Weighted networks, e.g., to include interaction confidence scores, are supported. (B) Supervised drug combination prediction with XGB using the features from (A) and ground truth data including preprocessing steps (feature selection, data balancing) and hyperparameter tuning.

### Network-based feature generation

Network-based features for CombDNF are generated using protein-protein interaction (PPI) networks. The proximity of specific nodes, e.g., disease nodes or drug targets, in PPI networks has been shown to be influential in diseases as well as for single drug and drug combination prediction (Guney et al., 2016; Cheng et al., 2019). We therefore focus on those features that characterize the mode of action of a drug in the context of a disease and an organism as input to CombDNF. Therefore, similar to Cheng et al. (2019), for a disease and a combination of two drugs, we consider three sets of nodes or proteins in the PPI network and their distances: the drug targets of each of the drugs, and a set of proteins encoded by genes relevant for the disease in question (disease genes). As features, several disease- and drug-specific distance measures are computed based on shortest paths in the PPI network, and on further node set characteristics.

### PPI network, disease genes, drug targets

The Human Integrated Protein–Protein Interaction rEference (HIPPIE) v2.3 (Alanis-Lobato et al., 2016) is used as a weighted protein-protein interaction network. We choose HIPPIE as a regularly updated PPI network with high coverage of proteins and manually curated, high-quality interactions. In addition, it provides confidence scores of the protein interactions as additional information that enables weighting the network edges by their respective confidence (see Supplementary Note 1 for details). The largest connected component of the network that we use consists of 19,669 proteins (nodes) and 827,106 interactions (edges).

Disease-specific features are derived by computing distances to the set of disease genes, i.e., disease-specific proteins that are associated with the disease in question. The Comparative Toxicogenomics Database (CTD, version Jan 31, 24 (Davis et al., 2022)) is queried based on Medical Subject Headings (MeSH) IDs to extract sets of disease genes for each of the four diseases. Only genes with direct evidence are considered (see Supplementary Note 1 for details and numbers of disease genes per disease). We also investigate disease genes from the Precision Medicine Knowledge Graph (PrimeKG) database (Chandak et al., 2023) as an alternative. However, this only has minor effects on the prediction performance (see Supplementary Note 6 for details).

Drug target information is extracted from Cheng et al. (2019) based on high-quality drug target interactions from various sources. A known molecular mode of action of a drug is essential for assessing drug interactions for predictions with CombDNF. Thus, drugs and their combinations are only included in the dataset if they have at least one known drug target appearing in the PPI network (see Supplementary Note 1 for the respective reduction in drugs per disease).

### Network-based distances as features

We leverage two distance-based scores that characterize drugs and diseases for network-based feature generation (see Supplementary Note 2 for details). We adapt both score computations from Cheng et al. (2019) to include edge weights to leverage more fine-grained information via PPI confidences. Thereby, we use the additive inverse of the interaction confidence scores plus one as edge weights to account for reduced confidence as increased cost of an edge in the shortest path computations. Then, we rely on the weighted version of the shortest path algorithm as the basis for distance computations. First, we compute the weighted version of the separation score *s*_*XY*_ (Cheng et al., 2019) as a symmetric measure of the mean shortest path between two sets of nodes *X, Y*. We employ it to characterize all pairwise distances between the two sets of drug targets and the set of disease genes. Second, for assessing the relationship between a drug and the disease, we calculate the weighted version of the Cheng et al. z-score of the closest distance between each set of drug targets and the disease genes. It measures the average shortest distance compared to the average shortest distances between random node sets with similar characteristics. As the Cheng et al. z-score is not symmetric, we use both orders of the arguments as two separate distance-based features per drug-disease pair.

In addition, we compute the number of overlapping proteins, as well as the mean, the median, the minimal, and the maximal (weighted) shortest path lengths between each pair of node sets as additional new features. Overall, we employ 22 features for each drug-drug combination and disease (see Supplementary Table S2 for a full list).

### ML-based classification with CombDNF

Our drug combination classification method CombDNF employs XGB to classify drug combinations based on the above described features. The pipeline to train and evaluate CombDNF includes optional data processing steps: data scaling, data sampling, and feature selection (see Supplementary Note 3 for details).

For cross-validation, the data is divided into train, validation, and test sets with a class-label stratified split. *k*-fold cross-validation is used for hyperparameter tuning and model validation, and *k* evaluation scores are obtained. Further, CombDNF delivers both class labels and prediction probabilities.

As the prediction task is characterized by class imbalance, we employ adequate metrics for this setting. In particular, we use Matthews correlation coefficient (MCC) and area under the precision-recall curve (AUPR) for evaluation. We also report the area under the receiver operator curve (AUROC) for better comparability with other approaches. The latter two are computed from the prediction probabilities for the positive class.

In all experiments, for training CombDNF, we use a 5-fold cross-validation with 60-20-20 train-validation-test sets and over-sampling with ADAptive SYNthetic sampling (ADASYN, He et al. (2008), see Supplementary Note 3 for other settings). Hyperparameter tuning is based on optimizing the MCC (see Supplementary Table S3 for all employed values).

To avoid data leakage that hampers generalization abilities of models (Lo and MacKinlay, 1990), for our CombDNF method, we rely on the following setup. (i) For sklearn’s (Pedregosa et al., 2011) group *k*-fold, each drug combination is defined as a group independent of the order of the drugs. Therefore, we can ensure that each drug combination is only in either the train set, the validation set, or the test set. (ii) All data processing is performed strictly only after the split into the train-validation-test sets for each fold separately. Also, any required adaptations are only trained on the train set of each fold, and they are only applied (not adapted) to the validation and test sets. (iii) Other hyperparameter tuning is restricted to optimizing the validation sets within the train-validation-test splitting of the *k*-fold cross-validation. Further, all *k* models are provided for new predictions. Thus, overfitting of the model architecture to the test sets is reduced, and a measure of robustness and variability is provided along with the new predictions.

### State-of-the-art methods

We compare CombDNF to all identified state-of-the-art methods that (i) provided code to their methods, (ii) are reproducibly described, and (iii) are applicable to at least one of our four diseases (see Table 1).

Cheng et al. (2019) proposed an unsupervised disease-specific classification that relies on unweighted network-based distance features as we employ them for CombDNF. These features give rise to six threshold-based, disease-specific exposure patterns for drug combinations. They interpreted two of these exposure patterns as effective and adverse drug combinations, respectively. For our evaluation, we compute and interpret the exposure patterns as described in Cheng et al. (2019) based on the binary HIPPIE network. In case a drug combination cannot be classified, its predictions are treated as NA in our evaluation.

Shan et al. (2023) employed a multi-relational drug-disease network to train their VGAETF (Variational Graph Autoencoder Tensor Decomposition) model for classifying drug-drug-disease triplets. They constructed the drug-disease network from drug-disease mapping via MeSH codes and disease-disease relations from literature. As no trained model was provided, we train the VGAETF model with the ground truth and further input data (e.g., the required drug-disease multi-relational graph) as described by the authors. We use the mean of the prediction values over their cross-validation folds for the evaluation of their predictions.

Federico et al. (2022) proposed a genetic algorithm to detect disease-specific drug combinations for five cancer subtypes. These were identified based on minimal similarity of the drugs’ mechanism of action (from LINCS signatures (Stathias et al., 2020)), chemical structures, target closeness, and coverage in a cancer network. The cancer network was built from disease-specific gene expression data and a knowledge graph. As this approach is applicable to cancer data only, we include it only for the neoplasms-specific analyses. We combine their predictions for the five cancer-subtypes (see Supplementary Note 5 for more details). To fairly compare these three methods with CombDNF, we create a common data set from our ground truth data for evaluation. For that, we remove all drug combinations occurring in the training data of VGAETF and all drugs that did not appear at least once in a combination in their training data, thus avoiding data leakage. For evaluation, we use the intersections of our cross-validation test folds with the common data set (see Supplementary Table S7 for quantities). For assessing the effects of CombDNF’s design choices, we also train CombDNF using network-based features from a binary, unweighted PPI as well as an all-disease CombDNF model by concatenating features and ground truth of all four diseases.

For the MCC, we use zero as the decision boundary for the predicted values of all competitor methods, as described for Cheng et al. (2019). AUPR and AUROC are computed with each methods’s specific predicted values, i.e., probabilities of CombDNF, separation scores of Cheng et al., prediction values of VGAETF, and ranks of Federico et al.

### Drug combination ground truth data

CombDNF relies on comprehensive disease-specific ground truth data for supervised training. Therefore, we collect ground truth data from different complementary sources providing clinically approved and validated drug combinations (effective drug combinations) as well as clinically observed adverse effects when drugs are administered together (adverse drug combinations), see Figure 2 for an overview. Effective drug combinations are retrieved from Das et al. (2019), DrugCombDB (Liu et al., 2020), Cheng et al. (2019), Continuous Drug Combination Database (CDCDB, Shtar et al. (2022)), and DrugBank (Wishart et al., 2018) (see details in Supplementary Figure S1). Adverse drug combinations are gathered from Cheng et al. (2019) and DrugBank (Wishart et al., 2018).

**Fig. 2:**
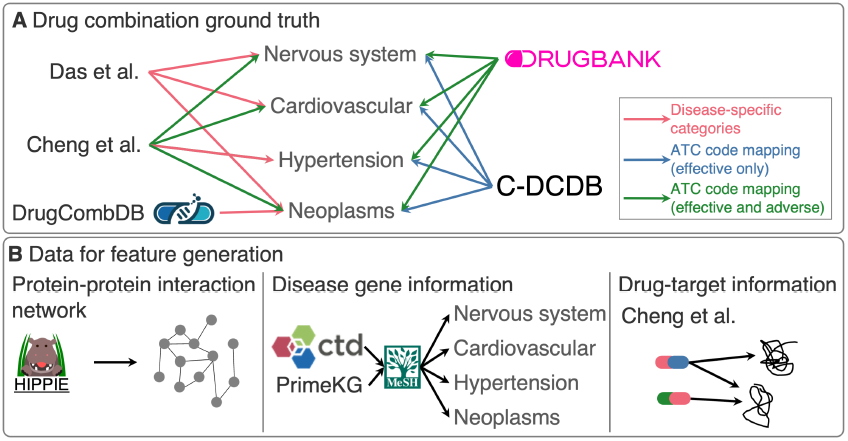
Overview of collected data for drug combination classification. (A) Disease-specific drug combination ground truth from clinically validated sources: Das et al., DrugCombDB, Cheng et al., CDCDB, and DrugBank. Arrows indicate the type of information that is contributed. (B) Data sources for disease-specific network-based feature generation for drug combinations: protein-protein interaction network from HIPPIE, disease-specific gene information (disease genes from CTD or PrimeKG), and drug-target information from Cheng et al.

As the effectiveness of drug combinations is by definition disease-specific, we also employ the ground truth combinations in a disease-specific manner. The assignment of a drug combination to a disease is either based on the anatomical therapeutic chemical (ATC) codes of contributing drugs, or on drug combination indication and literature information (see Figure 2 and Supplementary Note 1 for details, and Table 2 for quantification of the collected data; drug names are mapped to DrugBank IDs).

**Table 2.** Overview of drug combinations with clinically validated ground truth per disease. The number of effective and adverse drug combinations with ground truth that can be used for CombDNF is shown per disease. Both drugs in these combinations have drug targets within the PPI network. We indicate the number of unique drugs among the drug combinations, E/A as the effective-to-adverse ratio, and the density as the number of total ground truth combinations divided by the number of all possible combinations of unique drugs in each dataset.

| Disease | Drug combinations |  | E/A | Unique drugs | Density |
| --- | --- | --- | --- | --- | --- |
|  | Effective | Adverse |  |  |  |
| nervous system | 261 | 140,009 | 0.0019 | 1,400 | 0.1432 |
| cardiovascular | 279 | 85,545 | 0.0033 | 1,388 | 0.0892 |
| hypertension | 78 | 11,833 | 0.0066 | 1,154 | 0.0179 |
| neoplasms | 570 | 33,613 | 0.0170 | 1,330 | 0.0387 |

## Results

### Comparison to state-of-the-art prediction methods

First, we perform a benchmark analysis of disease-specific drug combination predictions with our novel method CombDNF. Therein, we compare its performance in four diseases with different characteristics with all available state-of-the-art methods identified in Table 1. In addition, using our model setup, we assess the influence of using unweighted instead of weighted network-based features, and the impact of single-disease-specific training vs. training one model of all diseases combined (see Table 3, and Supplementary Figure S4 for corresponding boxplots). All methods are evaluated on the same test dataset (over five cross-validation folds, see Methods for details). Our CombDNF models outperform all state-of-the-art methods in all performance scores and for all diseases (Federico et al. available for neoplasms only). Notably, while the competitor methods show acceptable accuracies, their prediction quality when evaluated by MCC and AUPR as is appropriate for label-imbalanced settings (Chicco and Jurman, 2020) is low. Further, we find CombDNF with unweighted network-based features (as used by Cheng et al. (2019)) also performing better than Cheng et al. (2019) on all datasets and all evaluation metrics. This substantiates that for the present prediction task, introducing supervised learning proves useful

**Table 3.** Performance comparison of drug combination classification for four diseases. CombDNF is compared to the approaches by Cheng et al. (2019), VGAETF (Shan et al., 2023), and Federico et al. (2022) (neoplasms only) on the same test sets (see Methods). We also assess CombDNF with features from unweighted, binary PPI networks (CombDNF unw.), and a single CombDNF model on data from all four diseases together (CombDNF all-d.). Mean performance scores are indicated with respective standard deviations over the five cross-validation folds (best mean scores per disease in bold). Our method CombDNF performs better than all state-of-the-art methods for all evaluation scores and diseases. Including edge weights for the network-based features improves predictions, and for 3/4 diseases, the single-disease CombDNF models perform better than the all-disease version. ^*^ Same test set as for the other methods but with different folds.

| method | MCC | AUPR | AUROC |
| --- | --- | --- | --- |
| **nervous system** |  |  |  |
| Cheng et al. | -0.02 ± 0.062 | 0.006 ± 0.002 | 0.433 ± 0.164 |
| Federico et al. | — | — | — |
| VGAETF | 0.003 ± 0.021 | 0.008 ± 0.004 | 0.496 ± 0.077 |
| CombDNF (ours) | **0.07** ± 0.064 | 0.045 ± 0.038 | 0.777 ± 0.056 |
| CombDNF (unw.) | 0.034 ± 0.061 | 0.039 ± 0.034 | 0.733 ± 0.063 |
| CombDNF (all-d.)* | **0.07** ± 0.157 | **0.097** ± 0.09 | **0.865** ± 0.144 |
| **cardiovascular** |  |  |  |
| Cheng et al. | -0.024 ± 0.026 | 0.01 ± 0.006 | 0.414 ± 0.077 |
| Federico et al. | — | — | — |
| VGAETF | -0.007 ± 0.001 | 0.011 ± 0.004 | 0.521 ± 0.061 |
| CombDNF (ours) | **0.124** ± 0.048 | **0.068** ± 0.013 | **0.754** ± 0.034 |
| CombDNF (unw.) | 0.057 ± 0.058 | 0.039 ± 0.023 | 0.684 ± 0.065 |
| CombDNF (all-d.)* | -0.002 ± 0.003 | 0.026 ± 0.01 | 0.657 ± 0.068 |
| **hypertension** |  |  |  |
| Cheng et al. | 0.049 ± 0.071 | 0.035 ± 0.023 | 0.57 ± 0.217 |
| Federico et al. | — | — | — |
| VGAETF | -0.013 ± 0.003 | 0.04 ± 0.016 | 0.589 ± 0.105 |
| CombDNF (ours) | **0.618** ± 0.225 | **0.665** ± 0.275 | **0.92** ± 0.078 |
| CombDNF (unw.) | 0.538 ± 0.227 | 0.52 ± 0.272 | 0.897 ± 0.097 |
| CombDNF (all-d.)* | 0.601 ± 0.383 | **0.665** ± 0.26 | 0.89 ± 0.133 |
| **neoplasms** |  |  |  |
| Cheng et al. | -0.035 ± 0.067 | 0.02 ± 0.003 | 0.461 ± 0.079 |
| Federico et al. | -0.009 ± 0.003 | 0.016 ± 0.004 | 0.497 ± 0.001 |
| VGAETF | -0.011 ± 0.012 | 0.02 ± 0.003 | 0.529 ± 0.048 |
| CombDNF (ours) | **0.575** ± 0.022 | **0.553** ± 0.047 | 0.884 ± 0.013 |
| CombDNF (unw.) | 0.548 ± 0.043 | 0.514 ± 0.047 | **0.888** ± 0.02 |
| CombDNF (all-d.)* | 0.258 ± 0.249 | 0.337 ± 0.229 | 0.877 ± 0.06 |

When comparing our different CombDNF versions, we find that omitting network edge weights during feature computation results in a decrease in performance compared to using the weighted network-based features. We conclude that incorporating further information such as confidence scores as weights into the prediction is beneficial. Further investigations also show that our additionally introduced features (see Methods) are relevant as the prediction performance of CombDNF drops when restricting it to the features by Cheng et al. only, yielding, e.g., an average MCC of 0.23 compared to 0.66 for neoplasms (see Supplementary Table S9).

Further, our single-disease-specific models outperform our all-disease model for all diseases except for nervous system for all scores and hypertension for AUPR. We compare novel predictions of CombDNF for drug combinations that are shared between the diseases (see Supplementary Figure S9). Of note, we find that the predictions of our all-disease model are decisively more similar between diseases than the predictions of our single-disease CombDNF models. This indicates that the single-disease approach might be better able to represent specific characteristics of a disease during drug combination prediction.

### Effect of ground truth data characteristics

Next, we investigate a potential underlying reason for varying prediction quality of CombDNF between diseases: It might be attributed to differences in the characteristics of the underlying ground truth data. In particular, there is a higher ratio of effective-to-adverse drug combinations for neoplasms and hypertension than for the other two diseases (see Table 2).

To quantify the effect of label imbalance, we train our disease-specific CombDNF model on subsets of the ground truth dataset with different effective-to-adverse ratios (see Figure 3 for hypertension and neoplasms drug combinations, and Supplement Figure S6 for cardiovascular and nervous system). These subsets are created by successively removing all combinations of drugs with the lowest effective-to-adverse ratio. Thereby, increasing this ratio shows overall improved performances of CombDNF. This effect is less pronounced for the other two diseases, where we also achieve only smaller maximal effective-to-adverse ratios.

**Fig. 3:**
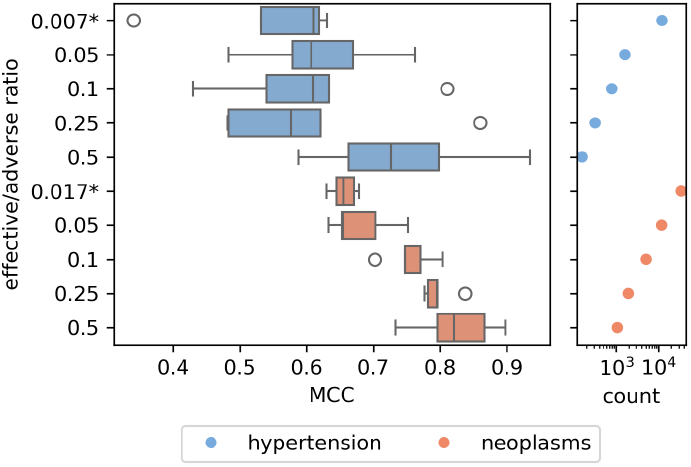
The effect of ground truth class label ratios for hypertension and neoplasms. The performance of CombDNF (MCC, left) trained on different subsets of the ground truth data with increasing effective-to-adverse ratios. Also the number of drug combinations (count, log-scale, right) in the data subsets is given. The effective-to-adverse ratio has a strong influence on the prediction, with worse overall performance in more imbalanced settings. * complete ground truth dataset compared to the unsupervised threshold-based drug combination exposure configuration definition by Cheng et al. (2019).

The same mechanism could be the reason why the all-disease CombDNF performs better than the single-disease model for nervous system diseases in our case (effective-to-adverse ratio 0.0019 for single-disease vs. 0.0043 for the all-disease setting, see also Table 3). Consequently, in highly label-imbalanced settings, an all-disease model may be beneficial.

We observe similar results when comparing CombDNF trained on single vs. the combined ground truth data sources (see Supplementary Note 6). In 3/4 diseases, performance is best when training and evaluating on the data source with the highest effective-to-adverse ratio, but these datasets are smaller, effectively reducing the number of drugs for which predictions can be made. In contrast, while also variable between disease datasets, the density of the ground truth datasets does not have a strong effect on prediction performance (see Supplementary Note 6).

### Biological relevance of predictions by CombDNF

We investigate potentially effective (top ten scoring) new drug combinations for neoplasms-specific CombDNF for cancer treatment (see Supplementary Table S10-S13 for effective and adverse drug combination predictions for all diseases). Indeed, we find evidence that the top combinations predicted by CombDNF are biologically plausible. First, the highest-scoring drug combination contains the anti-neoplastic drug Cediranib together with Vatalanib. Both drugs are mainly vascular endothelial growth factor receptor (VEGFR) inhibitors. However, Wissner et al. (2007) found that additionally to VEGFRs, Vatalanib also inhibits the epidermal growth factor receptor (EGFR). Studies showed that combining VEGFR inhibitors with inhibitors of other receptor tyrosine kinases including EGFR can be beneficial in treating different types of cancer (Liu et al., 2022a). Therefore, we see evidence that combining Cediranib with Vatalanib could be effective for cancer treatment. The simultaneous inhibition of VEGFRs and EGFR is also a possible underlying mechanism of action of two additional of our top predicted drug combinations: Vatalanib in combination with the VEGFR-inhibitor Bevacizumab (score: 0.989), and Cediranib together with Canertinib, an EGFR and pan-erbB tyrosine kinase inhibitor (score: 0.972).

**Table 4.**
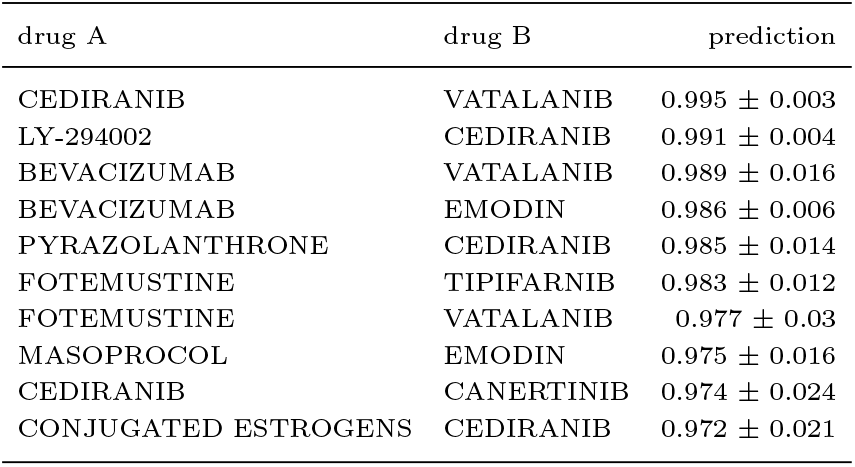
Top ten scoring potentially effective new drug combination predictions of CombDNF for neoplasms. We only consider combinations containing at least one cancer-specific drug (ATC code L01). Mean and standard deviation of prediction probabilities over the five-fold cross-validation CombDNF models are displayed.

## Discussion and Conclusion

We here propose a novel method for disease-specific prediction of clinically relevant drug-drug interactions, CombDNF. We conduct performance benchmarking in four diseases with diverse characteristics. Here, our approach CombDNF outperforms the current state-of-the-art methods across all diseases, improving AUPR scores by at least 84%. This strong improvement in performance is likely due to training an ML model also on clinically reported adverse combinations. Further reason for improvement are network-based features representing the molecular basis of the disease and drug action. Furthermore, predictions improve by explicitly representing uncertainty associated with interactions in the employed PPI network by including interaction confidence scores for the edge weights. We find evidence for the predictions of CombDNF to also be biologically meaningful. For example, it predicts cancer drug combinations as effective that target the complementary VEGF and EGF receptors, a mechanism that has been described in the literature as beneficial for cancer treatment (Liu et al., 2022a). In addition, the disease-specific modeling approach of CombDNF commonly provides an advantage over a generic all-disease approach. We find that prediction performance is strongly influenced by the effective-to-adverse ratio in the ground truth data, with models having difficulties providing acceptable prediction performance in highly imbalanced settings. In these cases, choosing more selective databases for training and prediction could be helpful although these often have the disadvantage of a reduced number of drugs for which predictions can be derived. Alternatively, the all-disease approach that enables CombDNF to learn from additional data could be beneficial.

For CombDNF’s training and evaluation pipeline, we put emphasis on reproducibility and avoiding data leakage. This is a common problem hindering the application and generalization ability of published models (Lo and MacKinlay, 1990; Bernett et al., 2024). Consequently, for evaluation of VGAETF and our CombDNF all-disease version, we did not allow the drug combination of a drug-drug-disease triplet to appear in both the train and the test set. This, in addition to the strong label imbalance, could be a reason for the observed differences to the performance measures reported in the original publication of VGAETF (Shan et al., 2023).

Due to limited availability of clinically validated ground truth data, our models are trained and evaluated in a warm-start scenario. This means that our estimated prediction quality is only representative for the classification of drug combinations where drugs appear at least once (in combination with other drugs) in the training set (Liu et al., 2022b). Stratifying the data splits by excluding all combinations of a drug in the train or validation set could extend the application range of CombDNF to predictions for combinations including unseen drugs, but this would require more training data as this is a more difficult prediction task. While CombDNF can make predictions for all provided combinations, a warning is issued for those drug combinations without any ground truth in the train set.

A limiting factor of CombDNF is its dependence on known high-quality drug targets which reduces the number of drug combinations for which CombDNF can derive predictions (see Supplementary Table S1). While this enables including valuable molecular underpinnings of drug action, it might also result in potential biases of drugs that can be predicted with our method (see Supplementary Note 1 for ATC code distribution). Neoplasms drug combinations are among the most affected by this reduction, but CombDNF’s predictions are good for the remaining drugs for neoplasms.

Instead of HIPPIE as a highly reliable, curated PPI network that CombDNF uses for its disease-specific network-based feature generation, other graphs could be employed. Examples are STRING (Szklarczyk et al., 2023), or ConsensusPathDB (Kamburov and Herwig, 2022). In particular, a directed protein- (or gene-)regulatory network (Liu et al., 2015) could add more fine-grained information on cellular function to features of CombDNF and could be worthwhile exploring.

Apart from other molecular interaction networks, CombDNF might benefit from including disease-specific information in the structure of its (currently disease-agnostic) PPI network, an avenue that should be explored in the future. If successful, changes to the PPI network of CombDNF could even be used to encode patient-specificity, e.g., from mutation (Löwer et al., 2012; Tate et al., 2019) or expression data (Moreno et al., 2022). This makes predictions more personalized but also reduces its application range as observed for Guo et al. (2021). However, since very limited clinical ground truth data is currently available on a single patient level (Madani Tonekaboni et al., 2018; Wu et al., 2022), training a truly personalized model for multiple drug combinations is, in our opinion, not feasible at the moment. A possible solution could be transfer learning with pre-training models on disease groups and fine-tuning to the disease subtype level. Alternatively, pre-training on synthetic data that recapitulates molecular mechanisms in simulation-based transfer learning approaches might be another viable step towards further personalization (Kleissl et al., 2023).

We provide disease-specific ground truth data and a novel method, CombDNF, for disease-specific drug combination classification improving the AUPR by at least 84% compared to the current state-of-the-art. This translates, for example, to four effective drug combinations among the top ten hypertension-specific predictions by CombDNF compared to one for the best competing method (see Supplementary Table S8). CombDNF can easily be extended to predictions in other diseases based on clinically validated data. We make all code, including a flexible benchmarking pipeline for training and evaluation, freely available.

