## Supplementary Information for "CombDNF: Disease-specific drug combination predictions with network-based features on clinically validated data"

2025

Contact:

**7 Supplementary Note 7: Predictions of new drug combinations 17**

### 1 Supplementary Note 1: Datasets and characteristics

#### 1.1 PPI network

For network-based features generation, we use the Human Integrated Protein–Protein Interaction rEference (HIPPIE) v2.3 (Alanis-Lobato et al., 2016) as a weighted protein-protein interaction (PPI) network. We decided for HIPPIE because it is a regularly updated PPI network with high coverage of proteins and manually curated, high-quality interactions. In addition, it provides confidence scores of the protein interactions (Alanis-Lobato et al., 2016) that we use as valuable information for weighting the network edges (see Supplement for details). In order to enable the computation of reasonable distances as features, the network is reduced to its largest connected component consisting of 19,669 proteins (network nodes) and 827,106 interactions (network edges). As an extension to the approach by Cheng et al. (2019), we leverage the confidence scores of the PPIs and computed distances in the weighted PPI network. For that purpose, confidence scores are converted (1 - confidence) and used as edge weights, thus ensuring shortest paths via high-confidence connections.

#### 1.2 Disease genes

The Comparative Toxicogenomics Database (CTD) (release date Jan 31, 24 Davis et al. (2022)) was queried to extract sets of disease genes for each of the four diseases from the disease-gene associations. Medical Subject Headings (MeSH) IDs are used to assign genes to specific diseases. The MeSH tree was downloaded from the NIH website (`https://www.nlm.nih.gov/databases/download/mesh.html`, accessed: Feb. 15, 2024) and used to extract all IDs of the sub-diseases for the above-mentioned diseases (MESH sub-tree roots: C04 (neoplasm), C10 (nervous system), C14 (cardiovascular), C14.907.489 (hypertension)). All genes associated to a MESH sub-tree root or to the ID of any of its sub-diseases were considered as disease genes for the disease in question.

An alternative set of disease genes for each of the four diseases was extracted from the Precision Medicine Knowledge Graph (PrimeKG) database (release date Apr 25, 2022, Chandak et al. (2023)) (see also S1). Here, all protein-disease associations were considered based on MESH disease and disease sub-type IDs and names extracted from MESH as described above.

The choice of the disease gene set for the network-based feature generation has only minor effects on the prediction performance (see Supplementary Figure S5).

**Table S1:** Description of data. Number of disease genes from CTD (Davis et al., 2022) or PrimeKG (Chandak et al., 2023) with nodes in the HIPPIE PPI (Alanis-Lobato et al., 2016) for each disease. Values in brackets correspond to respective numbers extracted from databases without reduction to genes overlapping with the PPI network. Number of approved and adverse drug combinations in the ground truth data with drug targets in the PPI network for each disease. Values in brackets show numbers extracted from respective databases in total without reducing to drugs with drug targets in the PPI network.

| | **Disease genes** | | **Drug combinations** | | **Unique drugs** |
|---|---|---|---|---|---|
| **Disease** | CTD | PrimeKG | approved | adverse | |
| nervous system | 2,291 (2,429) | 1,621 (1,671) | 261 (544) | 140,009 (464,674) | 1,400 (4,186) |
| cardiovascular | 1,449 (1,624) | 972 (1,005) | 279 (657) | 85,545 (287,715) | 1,388 (4,186) |
| hypertension | 241 (251) | 16 (16) | 78 (98) | 11,833 (38,289) | 1,154 (2,890) |
| neoplasms | 3,636 (3,937) | 1,765 (1,816) | 570 (2,031) | 33,613 (171,992) | 1,330 (4,183) |

#### 1.3 Drug combination ground truth

Our disease-specific drug combination ground truth for the classification task includes data from Das et al. (2019), DrugCombDB (Liu et al., 2020), Cheng et al. (2019), Continuous Drug Combination Database (CD-CDB) (Shtar et al., 2022), and DrugBank (Wishart et al., 2018) (see Figure 2, Supplementary Table S1, and their overlaps in Supplementary Figure S1).

Effective drug combinations were collected from multiple sources. We used drug combinations for nervous system diseases and cardiovascular diseases that are FDA-approved for the treatment of the respective diseases from Das et al. (2019). For neoplasms, in addition to FDA-approved drug combinations, also combinations that are clinically applied for cancer treatment from Das et al. (2019) were used. DrugCombDB provides a list of FDA-approved anti-cancer (neoplasm) drug combinations (release date May 31, 2019) (Liu et al., 2020). Cheng et al. (2019) assembled and published a list of non-disease specific FDA-approved and experimentally validated drug combinations from the literature in their Supplementary tables. They also provide a disease-specific list of combinations against hypertension curated from literature, which we used in addition to the combinations mapped via anatomical therapeutic chemical (ATC) codes as described below. The CDCDB compiles drug combinations from the FDA Orange Book (FDA-approved) with drug combinations from clinical trials and patents (release date June 18, 2024, Shtar et al. (2022)). Any combination that was withdrawn, terminated, or suspended in clinical trials and patents was removed. We consider all other combinations as effective and assigned them to the four diseases via ATC codes (see below). The DrugBank database (release 5.1.11) (Wishart et al., 2018) was queried for information on mixture products that have been marketed, i.e., used for treatment and containing two drugs that are both not withdrawn. This provides further effective combinations that were mapped to the investigated diseases via ATC codes.

Adverse drug-drug combinations were gathered from Cheng et al. (2019) and DrugBank (Wishart et al., 2018). Cheng et al. (2019) provide a list of clinically reported adverse drug combinations assembled from different sources. Additionally, we extracted all clinically reported adverse drug-drug interactions (collected from FDA drug labels and the literature) from the DrugBank database (Wishart et al., 2018), providing a list of adverse combinations.

If necessary, the drug names were mapped to their DrugBank ID. If the DrugBank ID was not found, the drug combination was not considered (2/157 and 69/946 drug combinations for Das et al. (2019) and DrugCombDB (Liu et al., 2020), respectively, were excluded).

All drug combinations, effective or adverse, from Cheng et al., CDCDB, and DrugBank were assigned to the four diseases using the anatomical therapeutic chemical (ATC) code information from DrugBank. Here, a drug combination was considered for a disease if at least one of the drugs is used to treat the disease according to ATC codes. We used ATC codes starting with L01 (neoplasm), N (nervous system), C (cardiovascular), and C02 (hypertension). While assembling the ground truth for each disease from all data sources, any duplicated combinations or combinations with conflicting labels were removed.

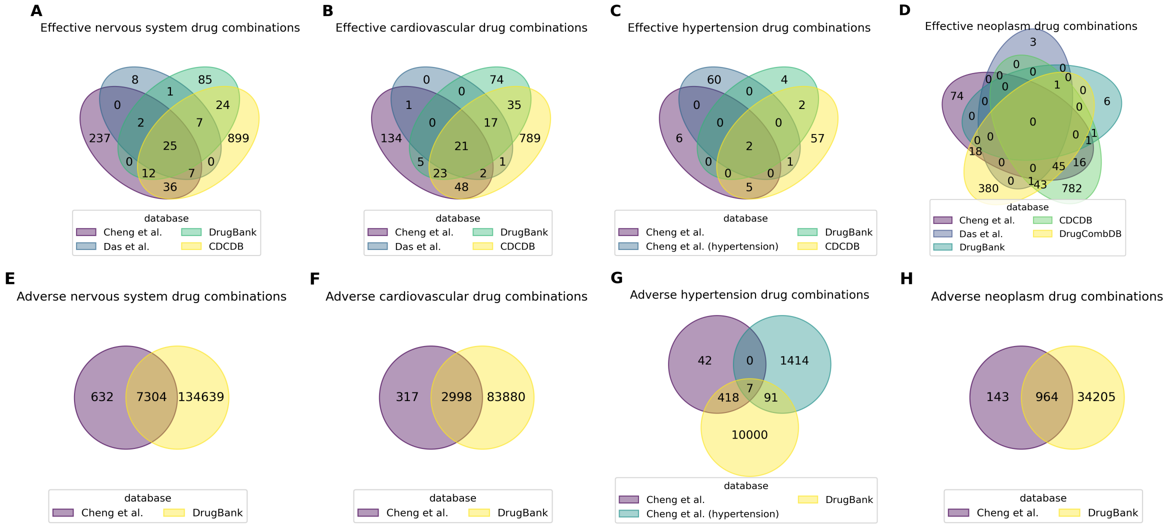

**Figure S1:** Drug combination from different data sources. (A)-(D) Respectively overlapping disease-specific effective drug combinations of drugs with target information and targets in our network from different data sources. (E)-(H) Respectively overlapping disease-specific adverse drug combinations of drugs with target information and targets in our network from different data sources.

#### 1.4 Drugs and drug combinations with targets in our PPI

Drugs and their combinations are only included in the ground truth dataset if they had at least one known drug target that was also contained as a node in the PPI network. This excluded up to 78% of drugs and up to 80% of drug combinations with ground truth data with slightly imbalanced distribution over the different disease groups (see Supplementary Table S1 and S2). While this requirement limits our considered ground truth dataset, a known mode of action of a drug is essential for assessing drug interactions and for making features disease-specific and derive disease-specific predictions with our approach. The distribution of the ATC codes shows that all disease groups are affected by a reduction in the number of drugs considered. Especially drugs without assigned ATC codes that cannot be considered for disease-specific predictions anyways are removed. Also for drugs with ATC codes J and V the decrease is considerable, for B, L (encoding neoplasms), P and A it is slightly less, see Supplement Figure S2.

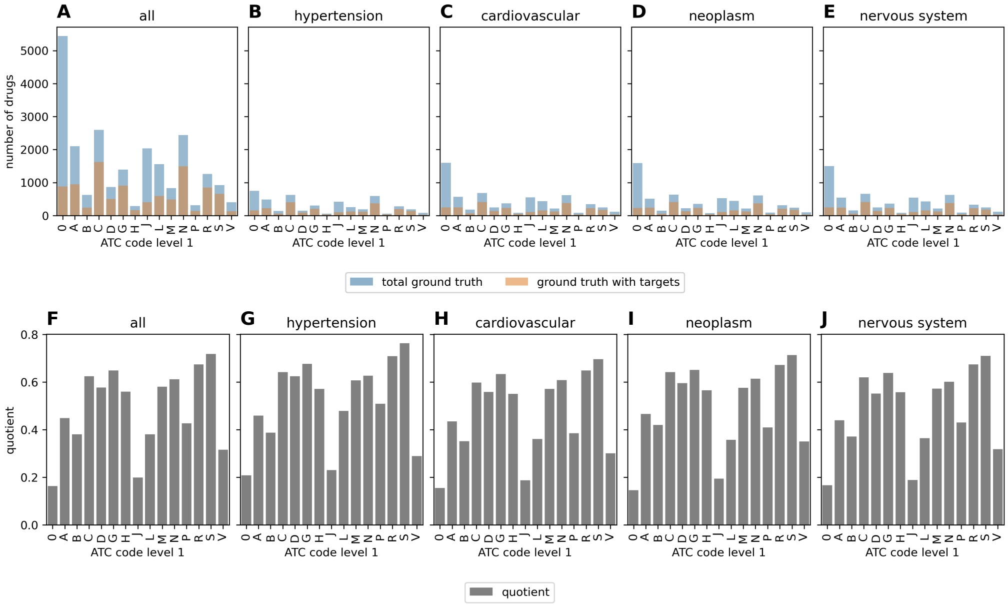

**Figure S2:** Distribution of Anatomical Therapeutic Chemical (ATC) codes. (A) Distribution of ATC codes of single drugs from all ground truth combinations (blue) for all four diseases combined and the distribution of ATC codes of drugs from drug combinations with targets in the PPI network (orange). (B)-(E) Distribution of ATC codes respectively for each of the four diseases. (F) The ratio between drugs with targets in the PPI network and all ground truth drugs divided by their ATC code. (G)-(J) Ratios respectively for each of the four diseases. Drugs with ATC code N are for nervous system disease, C for cardiovascular diseases, including hypertension, and L for neoplasms. Drugs with code 0 have no ATC code available in DrugBank (Wishart et al., 2018). Overall, drugs with all of the different ATC codes are affected by reduction when considering only those with drug targets in the PPI network. Especially, the without ATC codes (0), and with ATC codes J, V, B, L, P, and A are affected. For the other ATC codes, 50% or more of the drugs remain.

### 2 Supplementary Note 2: Network-based features

#### 2.1 Distance metrics (Cheng et al. (2019) scores) and features

We use two distance-based scores on our node sets (each drug's targets, set of disease genes) that characterize drugs and diseases for network-based feature generation that were shown to be useful for the task (Cheng et al., 2019). The scores were implemented by Cheng et al. (2019)for a binary, unweighted PPI network. We adapt both to include edge weights in the computation to leverage more fine-grained information via PPI confidences.

First, the separation score $s_{XY}$ from Cheng et al. as a symmetric measure of the mean shortest path between two sets of nodes $X$, $Y$ was used (see Supplement Eq. 1). Cheng et al. (2019) employed the separation score solely for measuring the distance between two drugs by comparing their sets of drug targets. For a fixed PPI network, this distance between drugs is disease-agnostic. In addition to that, we also use the separation score to assess the distance between a drug and a disease, using the drug's targets and the disease genes as node sets.

$$s_{XY} = \langle d_{XY} \rangle - \frac{\langle d_{XX} \rangle + \langle d_{YY} \rangle}{2} \tag{1}$$

where $\langle d_{XY} \rangle$, $\langle d_{XX} \rangle$, $\langle d_{YY} \rangle$ describe the mean shortest distance between the nodes of sets X and Y, X and X, Y and Y, respectively (Cheng et al., 2019).

Second, for assessing the relationship between a drug and the disease, we use the z-score from Cheng et al. of the closest distance, $z$, between the sets of drug targets and disease genes (Cheng et al., 2019) (see Supplement Eq. 2 and 3). It measures the average distance compared to average distances between random node sets with similar characteristics. This z-score is not symmetric between two sets of nodes, as averaging is only performed over one of the sets. Therefore, we use the z-scores of the closest distance, respectively, averaged over the drug targets of the considered drug ($zDT$) as well as averaged over the disease genes ($zTD$, also used in Cheng et al. (2019)) as two separate distance-based features per drug.

$$c(X,Y) = \frac{1}{||Y||} \sum_{y \in Y} \min_{x \in X} sp(x,y) \tag{2}$$

were $X$ and $Y$ are sets of nodes in the network, $||Y||$ is the number of nodes in set $Y$, and $sp(x,y)$ is the shortest path between nodes $x$ and $y$ (Cheng et al., 2019). The z-score $z$ is computed with the mean and standard deviation of the closest distance of 100 sets of random nodes with similar degree distributions:

$$z = \frac{c - \mu_{c_{100}}}{\sigma_{c_{100}}} \tag{3}$$

In addition, we compute the number of overlapping proteins, as well as the mean, the median, the minimal, and the maximal (weighted) shortest path lengths between each pair of node sets (drug targets-drug targets and drug targets-disease genes) as additional new features. Overall, 22 features are computed for each drug-drug combination and disease (see Supplementary Table S2 for a full list).

**Table S2:** Description of network-based features used for drug combination classification. Features that Cheng et al. (2019) use in their method for drug combination characterization are marked with *.

| **feature name** | **description** | **Cheng et al. features** |
|---|---|---|
| | **drug-drug features** | |
| sAB | separation score (Eq. 1) of drug targets of drug A & drug B | * |
| min_spAB | minimal shortest path between targets of drug A & drug B | |
| max_spAB | maximal shortest path between targets of drug A & drug B | |
| mean_spAB | mean shortest path between targets of drug A & drug B | |
| median_spAB | median shortest path between targets of drug A & drug B | |
| overlap_AB | number of proteins in common between targets of drug A & drug B | |
| | **drug-disease features** | |
| zTDA, zTDB | z-score (Eq. 3) of the closest distance (Eq. 2) for drug A/drug B averaged over the disease genes | * |
| zDTA, zDTB | z-score (Eq. 3) of the closest distance (Eq. 2) for drug A/drug B averaged over the drug targets | |
| sAD, sBD | separation score (Eq. 1) of targets of drug A/drug B and disease genes | |
| min_spAD, _spBD | minimal shortest path between targets of drug A/drug B and disease genes | |
| max_spAD, _spBD | maximal shortest path between targets of drug A/drug B and disease genes | |
| mean_spAD, _spBD | mean shortest path between targets of drug A/drug B and disease genes | |
| median_spAD, _spBD | median shortest path between targets of drug A/drug B and disease genes | |
| overlap_AD, _BD | number of proteins in common between targets of drug A/drug B and disease genes | |

### 3 Supplementary Note 3: Data balancing and feature selection and scaling in CombDNF

#### 3.1 Cross-validation

For cross-validation in CombDNF, the data is divided into train, validation, and test sets. The split is class-label stratified, i.e., class proportions are retained between the train, validation, and test sets. The parameter $k$ for $k$-fold cross-validation is a user-defined value. For each of the $k$ folds, the training and validation sets are used to train and tune CombDNF, respectively (see Supplementary Table S6 for a full list of hyperparameters). The test set in each fold is used to predict and evaluate the performance using the model trained on the train and validation set with the best hyperparameters. Thus, the pipeline provides $k$ evaluation scores, one for each of the $k$ test sets. Every data point is seen once during testing, which provides a more robust evaluation. Additionally, classes of new drug combinations, if provided by the user, can be predicted using the best model of each of the $k$ folds. Thus, for each new drug combination, $k$ predictions of $k$ models trained on slightly different data are obtained. This provides a measure of the variability of the predictions. In addition to the class predictions, probabilities of the predictions are provided for the $k$ test sets and new drug combinations, respectively.

#### 3.2 Optional data processing options

After data splitting, the pipeline of CombDNF includes different optional data processing steps: data scaling, data sampling, and feature selection. Scaling of the features is performed feature-wise with a z-score standardization. To balance the train set in highly class-imbalanced datasets, the following optional data sampling methods are included in CombDNF: ADAptive SYNthetic sampling (ADASYN, He et al. (2008)), Synthetic Minority Over-sampling Technique (SMOTE, Chawla et al. (2002)), SMOTE + Edited Nearest Neighbors (SMOTE+ENN, Batista et al. (2004)), and SMOTE + Tomek Links (SMOTE+Tomek, Batista et al. (2003)). The first two methods are oversampling methods, and the latter two combine over- and undersampling. Feature selection methods can help reduce the search space and may be beneficial for some of the ML methods. The pipeline supports two optional supervised feature selection methods: SelectKBest and recursive feature elimination (RFE, Guyon et al. (2002)) with Classification and Regression Trees (CART, Breiman et al. (1984)). SelectKBest selects the $k$ best features based on mutual information between the features and the ground truth. RFE with CART recursively eliminates features with low predictive importance based on a CART decision tree. Both feature selection methods reduce features to a user-defined number of features. To optimize the number of features, feature selection is included as a step in the hyperparameter tuning using 10% to 90% of the features. The data processing steps can be chosen by the user and are applied in the order described above. If required, data processing steps are trained only on the train set and are then used to transform the validation and test sets accordingly to avoid data leakage.

##### 3.2.1 Benchmark of data processing options

CombDNF natively allows for optimizing the choice of balancing approaches, feature scaling, and feature selection. We benchmark different data balancing methods, features selection methods, and scaling (see Supplementary Tables S4 and S5). For all diseases, we compute a baseline (no additional data processing) and a classification with ADASYN, SMOTE, SMOTE+ENN, and SMOTE+Tomek oversampling. Additionally, we compute classification with feature scaling and feature selection with $k$-best or RFE.

We find that each of the balancing methods, including oversampling, improves the predictions for all four diseases compared to the baseline, while feature selection and scaling alone show very little to no improvement in prediction performance (see Supplementary Table S5). Based on the MCC scoring that we consider most appropriate for this highly imbalanced setting, oversampling with ADASYN outperforms the other three methods for drug combination predictions in cardiovascular diseases, hypertension, and neoplasms (see Supplementary Table S4). For diseases of the nervous system, the over- and undersampling method SMOTEEN performs best (MCC of 0.07 compared to 0.05 with ADASYN).

**Table S3:** Hyperparameters and their values used for hyperparameter tuning of CombDNF.

| hyperparameters |
|---|
| `learning_rate: [0.001, 0.01, 0.1], n_estimators: [10, 100, 500], max_depth: [5, 10],`<br>`min_child_weight: [5, 10], gamma: [0.0, 0.2, 0.4], subsample: [0.5, 0.7, 0.9],`<br>`colsample_bytree: [0.5, 0.7, 0.9], reg_alpha: [0.0, 0.5, 1.0], reg_lambda: [0.0, 0.5, 1.0]` |

**Table S4:** Performance evaluation with MCC score of different data balancing/sampling methods for ComDNF and diseases (see Supplementary Note 3 for more information). The mean MCC score is given with respective standard deviation (best mean score per disease in bold). We analyzed a baseline (no additional data processing) and a classification with ADASYN, SMOTE, SMOTE+ENN, and SMOTE+Tomek oversampling. CombDNF with ADASYN oversampling outperforms all other methods for cardiovascular diseases, hypertension, and neoplasms. CombDNF with SMOTEEN over- and undersampling outperforms CombDNF with ADASYN for nervous system.

| **Disease** | **Baseline** | **ADASYN** | **SMOTE** | **SMOTEENN** | **SMOTETomek** |
|---|---|---|---|---|---|
| nervous system | 0.014 ± 0.031 | 0.053 ± 0.022 | 0.059 ± 0.032 | **0.069** ± 0.029 | 0.059 ± 0.032 |
| cardiovascular | 0.024 ± 0.033 | **0.101** ± 0.052 | 0.095 ± 0.038 | 0.082 ± 0.022 | 0.088 ± 0.039 |
| hypertension | 0.402 ± 0.126 | **0.547** ± 0.121 | 0.526 ± 0.153 | 0.531 ± 0.133 | 0.544 ± 0.165 |
| neoplasm | 0.537 ± 0.055 | **0.656** ± 0.02 | 0.655 ± 0.028 | 0.601 ± 0.037 | 0.648 ± 0.035 |

**Table S5:** Prediction performance with MCC score of feature selection methods RFE and k-best and standard scaling for different ML models and diseases. The mean MCC score is given with respective standard deviation (best mean score per disease in bold). We analyzed a baseline (no additional data processing) and classification with feature scaling and feature selection with $k$-best or RFE. Feature selection improves predictions for nervous system diseases, cardiovascular diseases, and hypertension compared to baseline, but performance decreases for neoplasm. Thus, feature selection and scaling show very little to no improvement in prediction performance

| **Disease** | **Baseline** | **Standard scaling** | **k-Best selection** | **RFE selection** |
|---|---|---|---|---|
| nervous system | 0.014 ± 0.031 | 0.014 ± 0.031 | **0.061** ± 0.085 | 0.047 ± 0.066 |
| cardiovascular | 0.024 ± 0.033 | 0.024 ± 0.033 | 0.026 ± 0.037 | **0.037** ± 0.059 |
| hypertension | 0.402 ± 0.126 | 0.402 ± 0.126 | 0.398 ± 0.171 | **0.412** ± 0.134 |
| neoplasm | **0.537** ± 0.055 | **0.537** ± 0.055 | 0.525 ± 0.047 | 0.516 ± 0.033 |

### 4 Supplementary Note 4: Benchmark of CombDNF with different ML models

We benchmark ComDNF against different ML methods: k-nearest neighbors (kNN), linear discriminant analysis (LDA), logistic regression (LogReg), naive Bayes (NB), random forest (RF), and support vector machine (SVM). Thus, we cover a wide range of ML methods with different concepts, e.g., linear and non-linear, distance-based, probabilistic, and tree-based approaches, and different model complexities. For each of the four diseases, a kNN, LDA, LogReg, NB, RF, and SVM model is trained. We use 5-fold cross-validation with the same stratified split for each of the ML models and CombDNF. All tested hyperparameters are shown in Supplementary Tables S3 and S6. The MCC was used for optimization during hyperparameter tuning.

We evaluated the prediction performance on the test sets with MCC, AUROC, and AUPR. Figure S3 shows MCC, AUPR, and AUROC performance with ADASYN oversampling. All performance scores show that CombDNF outperforms other classification methods for all diseases. This has been reported for several other prediction tasks before (Shwartz-Ziv and Armon, 2022; Liu et al., 2019). Random forest (RF) performs comparable or second best for all four diseases.

#### 4.1 Hyperparameter-settings for model training

**Table S6:** Hyperparameters and their values used for hyperparameter tuning ML methods for benchmarking against CombDNF. (kNN: k-nearest neighbors, LDA: linear discriminant analysis, LogReg: logistic regression, NB: naive Bayes, RF: random forest, and SVM: support vector machine)

| model | hyperparameters |
|---|---|
| kNN | `n_neighbors: [2, 5, 8, ..., 29], weights: [’uniform’, ’distance’], p: [1, 2],` |
| | `metric: [’minkowski’]+[’cosine’]` |
| LDA | `solver: [’svd’]+[’lsqr’, ’eigen’], shrinkage: [None, ’auto’, 0, 0.1, 0.2, ..., 0.9]` |
| LogReg | `C: [100, 10, 1, 0.1, 0.01], penalty: [’l1’, ’l2’], max_iter: [100000],` |
| | `solver: [’liblinear’, ’saga’]+[’lbfgs’, ’newton-cg’, ’newton-cholesky’, ’sag’],` |
| | `class_weight: [’balanced’, None]` |
| NB | `var_smoothing:` [$10^{-9}$, $10^{-8}$, ..., 10] |
| SVM | `C: [100, 10, 1, 0.1, 0.01], gamma: [’scale’, 10, 1, 0.1, 0.01], max_iter: [100000],` |
| | `kernel: [’rbf’, ’poly’, ’sigmoid’, ’linear’], class_weight: [’balanced’, None]` |
| RF | `n_estimators: [10, 100, 500], criterion: [’gini’, ’entropy’],` |
| | `max_features: [’sqrt’, None], max_depth: [5, 10, 50], min_samples_split: [2, 10],` |
| | `min_samples_leaf: [1, 3], bootstrap: [True, False], class_weight: [’balanced’, None]` |

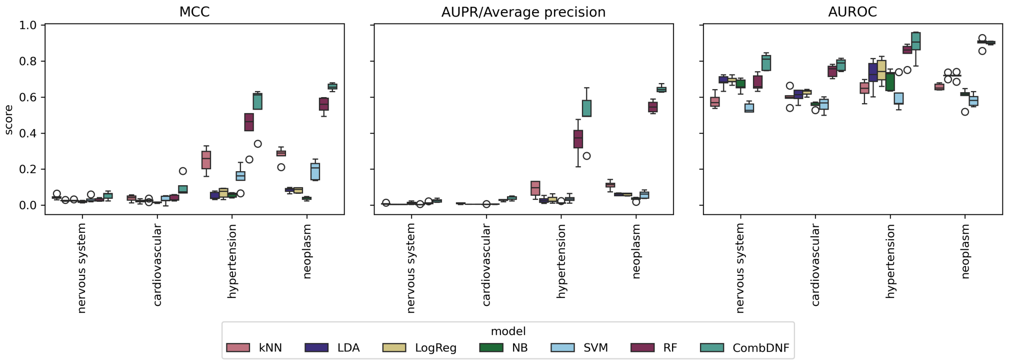

**Figure S3:** Benchmark of CombDNF and other ML methods for disease-specific drug combination classification with network-based features. Prediction performance is evaluated on (A) Matthews correlation coefficient, (B) area under the precision-recall curve (AUPR), and (C) area under the receiver operating characteristic curve (AUROC). Drug combinations were classified into adverse or efficient for four diseases with seven ML methods (see legend) using ADASYN oversampling (see Methods and Supplementary Note 4). The performance scores for the five cross-validation folds are shown as boxplots. For all diseases and evaluation metrics, CombDNF performed best or comparably with RF as runner-up. (kNN: k-nearest neighbors, LDA: linear discriminant analysis, LogReg: logistic regression, NB: naive Bayes, RF: random forest, and SVM: support vector machine)

### 5 Supplementary Note 5: Comparison with state-of-the-art methods

To fairly compare all state-of-the-art methods with CombDNF, we created a common test data set from our ground truth data for evaluation. For that, we remove all drug combinations that are in the train data of VGAETF (Shan et al., 2023) and all drugs that did not appear at least once in a combination in their training data, thus avoiding data leakage. This ensures all methods are evaluated on the same test sets. The methods from Chen et al. (2019) and Federico et al. (2022) do not provide predictions for all possible drug combinations with their methods, therefore predictions for these two were evaluated on smaller subet of the common test set (see Supplementary Table S7). For evaluation of all methods, we used the test sets from our cross-validation folds respectively reduced to combinations in the common data set (for amounts of ground truth data in the common test set, see Supplemental Table S7). Additionally, we compare CombDNF trained with features computed with the binary, unweighted PPI and with all diseases together (see below for Details). We compare prediction performance in Table 3 and Supplementary Figure S4. CombDNF outperforms all state-of-the-art methods for all diseases.

**Federico et al. (2022) predictions** Predicted drug combinations of Federico et al. (2022) were extracted from their Supplementary data for all five reported cancer types: breast cancer, liver hepatocellular carcinoma, prostate adenocarcinoma, colon adenocarcinoma, and lung squamous cell carcinoma. The drug combinations were ranked highest to lowest according to their occurrence in their solutions. Drug combinations with no occurrence were assigned the rank 0. Drug combinations from all five cancer types were combined, and the respectively highest rank was retained if duplicate drug combinations were identified. For prediction evaluation with MCC, drug combinations with rank one or higher were considered effective, and all others adverse. Performance evaluation with AUPR and AUROC is computed with the ranking.

**All-disease CombDNF** We investigated whether the all-disease model is beneficial in CombDNF and, therefore, trained CombDNF on data from all four diseases together (all-diseases model, see Methods). Note that no specific disease features were provided, but distance features were, in part, disease-specific. Overall, the all-disease model is based on a ground truth of 881,280 adverse drug combinations and 3,796 effective drug combinations, with an effective-to-adverse ratio of 0.0043. The prediction performance of the all-disease model is compared with those of the disease-specific models for each disease in Table 3 and Supplementary Figure S4. The all-disease model improves performance for drug combinations of nervous system diseases compared to the single-disease model but has a reduced performance for drug combinations of cardiovascular diseases, neoplasms, and hypertension. This suggests that predictions for diseases with lower effective-to-adverse ratios in their ground truth might benefit from an all-disease model.

**Table S7:** Number of effective and adverse drug combinations for the common test set. All drug combinations in our ground truth dataset are shown. Additionally, the number of combinations in the overlapping common test set for predictions with VGAETF (Shan et al., 2023), Cheng et al. (2019), and Federico et al. (2022) are given. For comparison with the SOTA methods the data set was reduced to a common test set (see Methods for details).

| Disease | Total ground truth | | Common ground truth test set with | | | | | |
|---|---|---|---|---|---|---|---|---|
| | | | VGAETF | | Cheng et al. (2019) | | Federico et al. (2022) | |
| | eff. | adv. | eff. | adv | eff. | adv. | eff. | adv. |
| nervous system | 261 | 140009 | 91 | 18623 | 20 | 3833 | - | - |
| cardiovascular | 279 | 85545 | 106 | 12909 | 36 | 4403 | - | - |
| hypertension | 78 | 11833 | 38 | 2043 | 20 | 1096 | - | - |
| neoplasm | 570 | 33613 | 213 | 12042 | 52 | 2681 | 38 | 2392 |

**Table S8:** Performance comparison of drug combinations classification with precision@$k$. The mean of precision@$k$ for the top five, ten, fifty, and hundred scoring drug combinations is given with respective standard deviation (best mean score per disease in bold). CombDNF outperforms state-of-the-art methods in correctly predicting the correct highest-scoring drug combinations. Especially for hypertension and neoplasm, CombDNF can predict correctly predict 50-100% of the highest-scoring drug combinations.

| Disease | Method | Precision@5 | Precision@10 | Precision@50 | Precision@100 |
|---|---|---|---|---|---|
| nervous system | Cheng et al. | 0.0 ± 0.0 | 0.0 ± 0.0 | 0.0 ± 0.0 | 0.0 ± 0.0 |
| | Federico et al. | nan ± nan | nan ± nan | nan ± nan | nan ± nan |
| | VGAETF | 0.0 ± 0.0 | 0.02 ± 0.045 | 0.004 ± 0.009 | 0.006 ± 0.009 |
| | CombDNF (our method) | 0.08 ± 0.11 | 0.1 ± 0.071 | **0.04** ± 0.028 | **0.034** ± 0.011 |
| | CombDNF (unweighted) | **0.12** ± 0.179 | **0.12** ± 0.13 | **0.04** ± 0.037 | 0.028 ± 0.022 |
| | CombDNF (all-disease) | 0.08 ± 0.11 | 0.04 ± 0.055 | 0.024 ± 0.009 | 0.02 ± 0.01 |
| cardiovascular | Cheng et al. | 0.0 ± 0.0 | 0.0 ± 0.0 | 0.008 ± 0.011 | 0.006 ± 0.009 |
| | Federico et al. | nan ± nan | nan ± nan | nan ± nan | nan ± nan |
| | VGAETF | 0.0 ± 0.0 | 0.0 ± 0.0 | 0.008 ± 0.018 | 0.012 ± 0.008 |
| | CombDNF (our method) | **0.24** ± 0.089 | **0.18** ± 0.084 | **0.072** ± 0.018 | **0.054** ± 0.023 |
| | CombDNF (unweighted) | 0.12 ± 0.179 | 0.1 ± 0.071 | 0.052 ± 0.054 | 0.036 ± 0.03 |
| | CombDNF (all-disease) | 0.0 ± 0.0 | 0.0 ± 0.0 | 0.016 ± 0.009 | 0.022 ± 0.011 |
| hypertension | Cheng et al. | 0.0 ± 0.0 | 0.02 ± 0.045 | 0.016 ± 0.022 | 0.024 ± 0.021 |
| | Federico et al. | nan ± nan | nan ± nan | nan ± nan | nan ± nan |
| | VGAETF | 0.0 ± 0.0 | 0.04 ± 0.055 | 0.028 ± 0.03 | 0.024 ± 0.023 |
| | CombDNF (our method) | **0.68** ± 0.179 | **0.46** ± 0.089 | **0.12** ± 0.024 | **0.062** ± 0.015 |
| | CombDNF (unweighted) | 0.56 ± 0.167 | 0.38 ± 0.084 | 0.108 ± 0.018 | 0.06 ± 0.012 |
| | CombDNF (all-disease) | 0.28 ± 0.11 | 0.16 ± 0.055 | 0.044 ± 0.017 | 0.028 ± 0.014 |
| neoplasm | Cheng et al. | 0.0 ± 0.0 | 0.0 ± 0.0 | 0.02 ± 0.02 | 0.014 ± 0.009 |
| | Federico et al. | 0.0 ± 0.0 | 0.02 ± 0.045 | 0.008 ± 0.011 | 0.01 ± 0.007 |
| | VGAETF | 0.0 ± 0.0 | 0.02 ± 0.045 | 0.008 ± 0.011 | 0.01 ± 0.01 |
| | CombDNF (our method) | **1.0** ± 0.0 | **1.0** ± 0.0 | **0.456** ± 0.074 | **0.264** ± 0.036 |
| | CombDNF (unweighted) | 0.96 ± 0.089 | 0.96 ± 0.055 | 0.448 ± 0.059 | 0.262 ± 0.04 |
| | CombDNF (all-disease) | 0.44 ± 0.329 | 0.3 ± 0.224 | 0.116 ± 0.033 | 0.072 ± 0.018 |

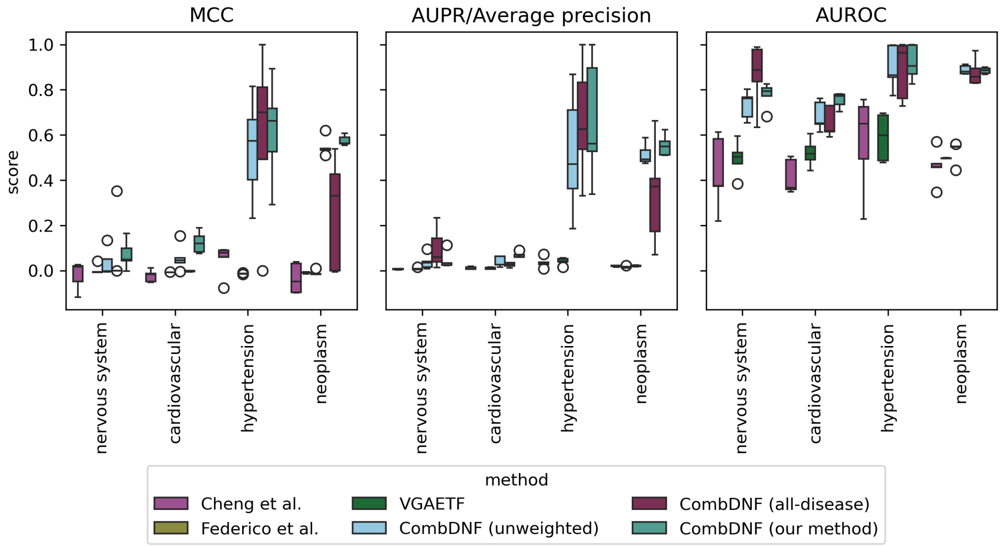

**Figure S4:** Comparison of state-of-the-art methods to our XGBoost model for drug combinations classification for four diseases (showing the data from Table 3). Prediction performance is evaluated on (A) Matthews correlation coefficient, (B) area under the precision-recall curve (AUPR), and (C) area under the receiver operating characteristic curve (AUROC). All five methods are evaluated on the same test set. The XGB (all-disease) model is evaluated on different cross-validation splits compared to the others.

### 6 Supplementary Note 6: Effect of ground truth data and features on prediction performance

#### 6.1 Effect of set of disease genes

We assess the effect of disease genes on the prediction performance of CombDNF using network-based features generated using disease genes from CTD (Davis et al., 2022) and PrimeKG (Chandak et al. (2023), see Supplementary Note 1 for more details). The choice of the disease gene set for the network-based feature generation has only minor effects on the prediction performance (see Supplementary Figure S5).

#### 6.2 Relevance of drug combination features

We also assess the relevance of the drug combination features for prediction quality. Thereby, we first investigate whether our drug combination features based on shortest paths in the weighted PPI network that we provide in addition to the ones used by Cheng et al. (2019) can be useful (see Methods and Supplemental Table S2 for a list of features). For this, we train CombDNF for all four diseases only on the features described by Cheng et al., derived from the weighted PPI network. We find that CombDNF with all features considerably outperforms CombDNF with only Cheng et al. features., yielding, e.g., an average MCC of 0.66 compared to 0.23 for neoplasms (see Supplementary Table S9).

In addition, we assessed the importance of particularly computationally demanding features proposed by Cheng et al., the two drug-disease z-scores, by training CombDNF without using these two features. The models without the computationally expensive features perform slightly worse than the model using all features (see Supplementary Table S9).

#### 6.3 Assessing varying ground truth data characteristics

To analyze the influence of ground truth data characteristics on the prediction performance of CombDNF, we create suitable subsets for training and evaluation. First, to investigate the effect of the effective-to-adverse ratio that varies between the collected ground truth of the different diseases (see Table 2), we create datasets with an increased effective-to-adverse ratio. We achieve this by successive removal of drugs and their combinations with the lowest effective-to-adverse ratio according to their ground truth (tested ratios 0.05, 0.1, 0.25, 0.5). CombDNF is trained and evaluated with the reduced ground truth data (see Supplementary Figure S6). Increasing the effective-to-adverse ratio shows improved performances for hypertension and neoplasms. This effect is less pronounced for the other two diseases, where we also achieved only lower efficient to adverse ratios up to 0.25.

Second, for investigating the influence of the density, i.e., the ratio of number of total ground truth combinations by number of all possible drug-drug combinations, on prediction performance, we create datasets with different increased density ratios. We achieve this by successive removal of drugs and their combinations with the lowest density ratio (tested densities 0.025, 0.05, 0.1, 0.25). We train and evaluate CombDNF on the reduced ground truth data with different data densities (see Supplementary Figure S7). For nervous system, cardiovascular, and hypertension drug combinations, we find only very few changes in performance even for drastically increased density, but the performance for neoplasm drug combinations drops with higher density. When inspecting the effective-to-adverse ratio for this experiment (middle of Figure Supplementary Figure S7), we find this ratio to be relatively stable for the first three diseases, while it is reduced with higher densities for neoplasm drug combinations. Thus, the performance drop could also be attributed to the reduced effective-to-adverse ratio in this case, and it seems to affect prediction performance to a greater extent than data density.

Third, we analyze the effect of different ground truth data sources on prediction performance. Therefore, for each database and each disease, we create a new (sub-)dataset only containing the database-specific drug combinations associated with the disease as described before. The neoplasm dataset from Das et al. (2019) is excluded from this analysis since it contains very few effective drug combinations S1. We use adverse combinations from DrugBank as the negative/adverse class for DrugCombDB (Liu et al., 2020) and CDCDB (Shtar et al., 2022) as these only capture effective combinations. As DrugCombDB is cancer-specific, we only analyzed it for neoplasms. In addition, Cheng et al. published a hypertension-specific ground truth with effective and adverse drug combinations, which is analyzed separately from the Cheng et al. ground truth that is based on mapping drug combinations to diseases via ATC codes. We assess the prediction performance of CombDNF with the different sources of ground truth data (see Supplementary Figure S8). The ground truth data sources differ in their effective-to-adverse ratios but also in the number of drug combinations and unique drugs. In line with our findings from before (Figure S6), except for neoplasms, the prediction performance is best when training and evaluating the data source with the highest effective-to-adverse ratio (the Cheng et al. datasets for nervous system and cardiovascular diseases, and the hypertension-specific Cheng et al. dataset). Therefore, the performance of the combined data sources is often even exceeded. However, in all three diseases, the improved performance comes at the cost of a decisively reduced number of drugs with ground truth for which the model can derive predictions (to less than half the drugs). Of note, for the full Cheng et al. dataset for hypertension, which is second-best in terms of effective-to-adverse ratio, prediction performance is poor, probably due to the extremely reduced amount of available data (only 11 effective combinations). This could be similarly the cause for the decreased observed performance for the neoplasms model when trained and evaluated on the Cheng et al. dataset compared to the combined data sources, despite the decisively higher effective-to-adverse ratio of the former. For neoplasms, only using DrugCombDB data yields the best performance, potentially because this database is focused on cancer medication.

Overall, we find the effective-to-adverse ratio to have a central influence on performance (more balanced is better), which comes at the expense of a reduced number of drugs for which predictions can be made.

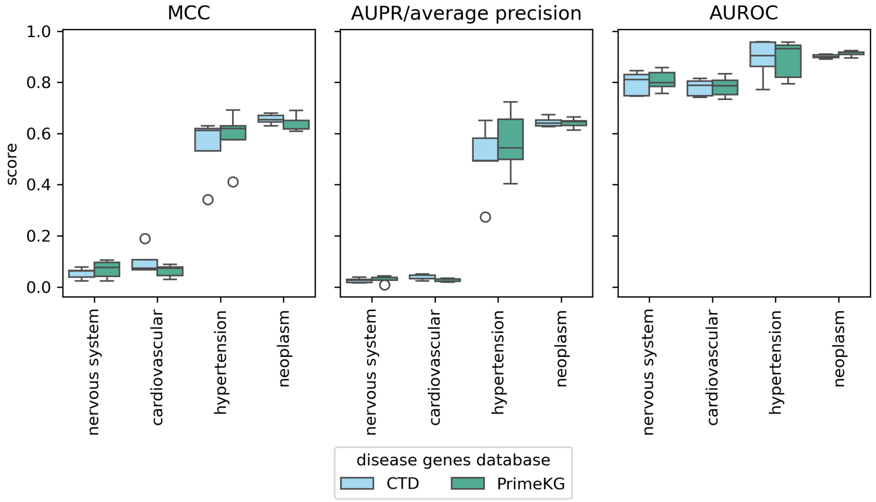

**Figure S5:** Effect of disease genes. Performance of network-based features from different disease gene databases with CombDNF for four diseases. Prediction performance is evaluated on: (A) Matthews correlation coefficient, (B) area under the precision-recall curve (AUPR), and (C) area under the receiver operating characteristic curve (AUROC). The databases of the disease genes are Comparative Toxicogenomics Database (CTD) (Davis et al., 2022) and Precision Medicine Knowledge Graph (PrimeKG) (Chandak et al., 2023). The choice of the disease gene set for the network-based feature generation has only minor effects on prediction performance.

**Table S9:** Feature influence on prediction performance. A comparison of the performance of CombDNF trained and tuned with original features from Chen et al. (2019), our features, and our features without the z-scores. The z-scores are computationally more expensive, as they require 100 randomized calculations. CombDNF with all features considerably outperforms CombDNF with only Cheng et al. features. CombDNF without the computationally expensive features performs slightly worse than the model using all features for almost all scores and diseases (except MCC for nervous system diseases).

| **Features** | **MCC** | **AUPR** | **AUROC** |
|---|---|---|---|
| | **nervous system** | | |
| Cheng et al. features | 0.028 ± 0.01 | 0.007 ± 0.002 | 0.684 ± 0.036 |
| our features | 0.053 ± 0.022 | **0.026** ± 0.009 | **0.796** ± 0.047 |
| our features without z-scores | **0.055** ± 0.029 | 0.021 ± 0.011 | 0.779 ± 0.028 |
| | **cardiovascular** | | |
| Cheng et al. features | 0.031 ± 0.007 | 0.008 ± 0.002 | 0.644 ± 0.036 |
| our features | **0.101** ± 0.052 | **0.037** ± 0.011 | **0.78** ± 0.033 |
| our features without z-scores | 0.076 ± 0.054 | 0.028 ± 0.01 | 0.742 ± 0.028 |
| | **hypertension** | | |
| Cheng et al. features | 0.195 ± 0.046 | 0.243 ± 0.118 | 0.848 ± 0.041 |
| our features | **0.547** ± 0.121 | **0.499** ± 0.142 | **0.891** ± 0.078 |
| our features without z-scores | 0.49 ± 0.173 | 0.492 ± 0.147 | 0.885 ± 0.083 |
| | **neoplasm** | | |
| Cheng et al. features | 0.225 ± 0.027 | 0.337 ± 0.055 | 0.798 ± 0.022 |
| our features | **0.656** ± 0.02 | **0.645** ± 0.019 | **0.901** ± 0.008 |
| our features without z-scores | 0.633 ± 0.04 | 0.607 ± 0.037 | 0.899 ± 0.012 |

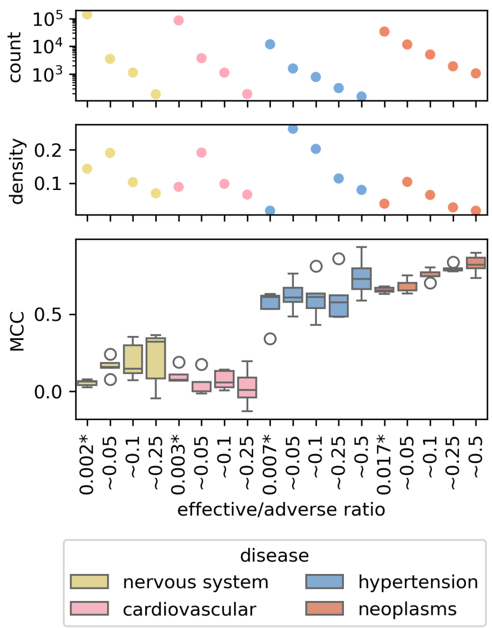

**Figure S6:** Prediction performance of CombDNF trained on ground truth data with varying class label ratios. The performance of CombDNF (MCC, bottom) trained on different subsets of the ground truth data with increasing effective-to-adverse ratios. Also, the density (middle) and number of drug combinations in the dataset (top) are given. * Unchanged ground truth.

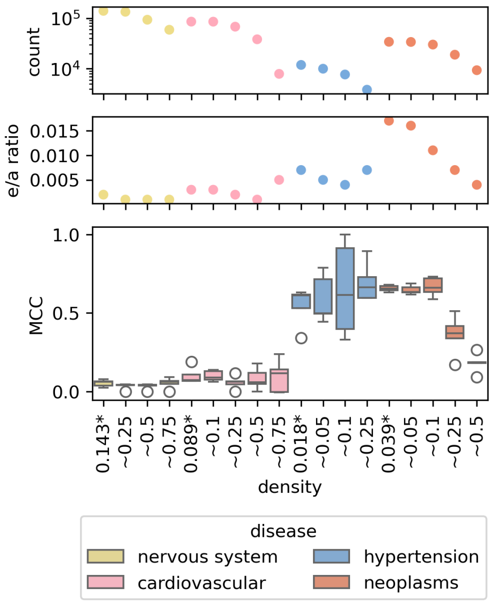

**Figure S7:** Prediction performance of CombDNF trained on ground truth data with varying data densities. The performance of CombDNF (MCC, bottom) trained on different subsets of the ground truth data with increasing density. Also, the effective-to-adverse ratio (middle) and number of drug combinations in the dataset (top) are given. * Unchanged ground truth.

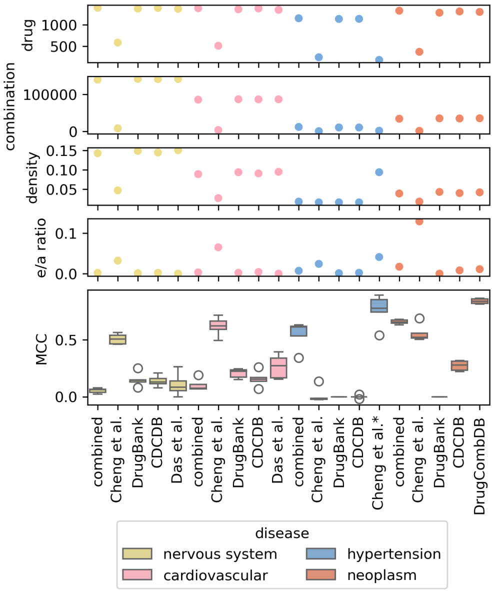

**Figure S8:** The effect of different ground truth sources for CombDNF performance. The CombDNF performance using different sources is compared with training on their combination (combined results as in Table 2). Along with the model performance (MCC, bottom), several dataset characteristics are shown (from top to bottom): number of unique drugs (drugs), number of drug combinations in the ground truth (combinations), density, and effective-to-adverse ratio (e/a ratio). We find the effective-to-adverse ratio has a central influence on performance (more balanced is better), which comes at the expense of a reduced number of drugs for which predictions can be made. * Hypertension-specific ground truth as provided by Cheng et al. (2019).

### 7 Supplementary Note 7: Predictions of new drug combinations

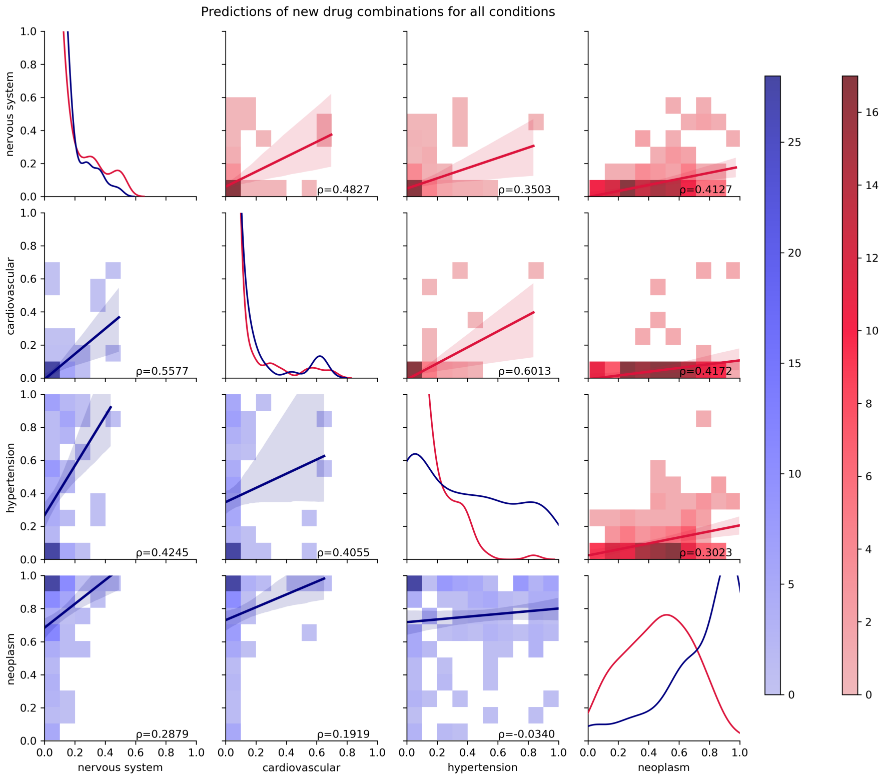

**Figure S9:** Correlation of ComDNF predictions for new drug combinations. We show predictions of 21 single drugs common in the effective combination of all four diseases and 206 drug combinations (drug combinations from ground truth are removed). A pairwise comparison of the mean of predicted probabilities over the five cross-validation models (close to 0 adverse; close to 1 predicted as effective in all five folds) is displayed with a linear regression with confidence interval and the Spearman correlation ($\rho$). The lower half of the matrix compares the predictions of the disease-specific CombDNF models. The upper half compares the predictions for the different diseases in the all-disease CombDNF model. On the diagonal, the distributions of predictions of the models are shown. Our single-disease model predictions have lower Spearman correlations between the diseases than the all-disease model (with the exception of cardiovascular and hypertension). This indicates that potential treatment differences can be better resolved using a disease-specific model. In particular, neoplasm predictions of the single-disease model have low correlations with the other three diseases. For nervous system and cardiovascular diseases, the distributions of predictions are very similar for the all-disease and single-disease models. For hypertension and neoplasm the single-disease predictions are shifted more towards 1 (effective).

**Table S10:** Top ten (potentially effective) and bottom ten (potentially adverse) predicted new drug combinations for neoplasms. Predictions are shown for drug combinations with at least one drug with an ATC code starting with L01 for neoplasm. The mean and standard deviation of prediction probabilities over the five-fold cross-validation models are displayed. Top predicted potentially effective drug combinations in neoplasms with anti-neoplastic drugs show prediction scores between 0.97 and 0.99 with small standard deviations of 0.03 or smaller over the models of the five folds. Thus, predictions of new drug combinations in neoplasms have considerably higher value and confidence than predictions in hypertension (see Supplementary Table S11).

| neoplasms | | | | |
|---|---|---|---|---|
| drugA name | drugB name | prediction probability | drugA ATC | drugB ATC |
| Top ten (potentially effective) drug combinations | | | | |
| CEDIRANIB | VATALANIB | 0.99487 ± 0.00266 | L01EK02 | NaN |
| LY-294002 | CEDIRANIB | 0.99112 ± 0.00412 | NaN | L01EK02 |
| BEVACIZUMAB | VATALANIB | 0.98854 ± 0.01573 | L01FG01; S01LA08 | NaN |
| BEVACIZUMAB | EMODIN | 0.98568 ± 0.00582 | L01FG01;S01LA08 | NaN |
| PYRAZOLANTHRONE | CEDIRANIB | 0.98493 ± 0.01413 | NaN | L01EK02 |
| FOTEMUSTINE | TIPIFARNIB | 0.98318 ± 0.01184 | L01AD05 | NaN |
| FOTEMUSTINE | VATALANIB | 0.97736 ± 0.02957 | L01AD05 | NaN |
| MASOPROCOL | EMODIN | 0.97497 ± 0.01624 | L01XX10 | NaN |
| CEDIRANIB | CANERTINIB | 0.97379 ± 0.02363 | L01EK02 | NaN |
| CONJUGATED ESTROGENS | CEDIRANIB | 0.97228 ± 0.02069 | G03CA57;G03CC07 | L01EK02 |
| Bottom ten (potentially adverse) drug combinations | | | | |
| TRETINOIN | PEMETREXED | 1e-05 ± 0.0 | L01XF01;D10AD51;D10AD01 | L01BA04 |
| CLOFARABINE | THIOTEPA | 1e-05 ± 1e-05 | L01BB06 | L01AC01 |
| TOPOTECAN | THIOTEPA | 1e-05 ± 1e-05 | L01CE01 | L01AC01 |
| MITOTANE | CLADRIBINE | 1e-05 ± 1e-05 | L01XX23 | L04AA40;L01BB04 |
| IRINOTECAN | REGORAFENIB | 1e-05 ± 1e-05 | L01CE02 | L01EX05 |
| VINCRISTINE | ANAGRELIDE | 1e-05 ± 0.0 | L01CA02 | L01XX35 |
| RITUXIMAB | THIOTEPA | 1e-05 ± 1e-05 | L01FA01 | L01AC01 |
| CABAZITAXEL | TRIMETREXATE | 1e-05 ± 0.0 | L01CD04 | P01AX07 |
| CABAZITAXEL | PIRITREXIM | 1e-05 ± 0.0 | L01CD04 | NaN |
| CABAZITAXEL | PROGUANIL | 1e-05 ± 0.0 | L01CD04 | P01BB01;P01BB51;P01BB52 |

**Table S11:** Top ten (potentially effective) and bottom ten (potentially adverse) predicted new drug combinations for hypertension. Predictions are shown for drug combinations with at least one drug with an ATC code starting with C02 for hypertension. The mean and standard deviation of prediction probabilities over the five-fold cross-validation models are displayed. Top potentially effective predictions in hypertension for anti-hypertensive drugs score between 0.65 and 0.78 with a relatively high standard deviation of 0.4. Thus, predictions of new drug combinations in neoplasms have considerably higher value and confidence (see Supplementary Table S10).

| hypertension | | | | |
|---|---|---|---|---|
| drugA name | drugB name | prediction probability | drugA ATC codes | drugB ATC codes |
| Top ten (potentially effective) drug combinations | | | | |
| ALPROSTADIL | MACITENTAN | 0.78309 ± 0.42858 | C01EA01;G04BE01 | C02KX04;C02KX54 |
| SITAXENTAN | ALPROSTADIL | 0.76973 ± 0.42006 | C02KX03 | C01EA01;G04BE01 |
| MINOXIDIL | FLOXURIDINE | 0.74222 ± 0.41705 | D11AX01;C02DC01 | L01BC09 |
| MINOXIDIL | PHENOLPHTHALEIN | 0.74222 ± 0.41705 | D11AX01;C02DC01 | A06AB04 |
| MINOXIDIL | CAPECITABINE | 0.74222 ± 0.41705 | D11AX01;C02DC01 | L01BC06 |
| MINOXIDIL | TRIFLURIDINE | 0.74222 ± 0.41705 | D11AX01;C02DC01 | L01BC59;S01AD02 |
| MINOXIDIL | FONDAPARINUX | 0.71554 ± 0.406 | D11AX01;C02DC01 | B01AX05 |
| MINOXIDIL | ENOXAPARIN | 0.71554 ± 0.406 | D11AX01;C02DC01 | B01AB05 |
| MINOXIDIL | HYPOXANTHINE | 0.65756 ± 0.43936 | D11AX01;C02DC01 | NaN |
| MINOXIDIL | DIDANOSINE | 0.65756 ± 0.43936 | D11AX01;C02DC01 | J05AF02 |
| Bottom ten (potentially adverse) drug combinations | | | | |
| CANAKINUMAB | MACITENTAN | 4e-05 ± 8e-05 | L04AC08 | C02KX04;C02KX54 |
| DROTRECOGIN ALFA | MACITENTAN | 4e-05 ± 8e-05 | B01AD10 | C02KX04;C02KX54 |
| DROTRECOGIN ALFA | BOSENTAN | 4e-05 ± 8e-05 | B01AD10 | G01AE10;C02KX01 |
| DROTRECOGIN ALFA | SITAXENTAN | 4e-05 ± 8e-05 | B01AD10 | C02KX03 |
| SITAXENTAN | TOLAZAMIDE | 4e-05 ± 7e-05 | C02KX03 | A10BB05;G01AE10 |
| SITAXENTAN | ACETOHEXAMIDE | 4e-05 ± 7e-05 | C02KX03 | G01AE10;A10BB31 |
| SITAXENTAN | TERAZOSIN | 4e-05 ± 7e-05 | C02KX03 | G04CA03 |
| METYROSINE | ACETOHEXAMIDE | 4e-05 ± 7e-05 | C02KB01 | G01AE10;A10BB31 |
| METYROSINE | TOLAZAMIDE | 4e-05 ± 7e-05 | C02KB01 | A10BB05;G01AE10 |
| CLONIDINE | GUANFACINE | 3e-05 ± 6e-05 | S01EA04;C02LC01;C02AC01;N02CX02;C02LC51 | C02AC02 |

**Table S12:** Top ten (potentially effective) and bottom ten (potentially adverse) predicted new drug combinations for cardiovascular diseases. Predictions are shown for drug combinations with at least one drug with an ATC code starting with C for cardiovascular diseases. The mean and standard deviation of prediction probabilities over the five-fold cross-validation models are displayed. The top potentially effective predictions in cardiovascular for anti-cardiovascular drugs score is between 0.65 and 0.78, with a relatively high standard deviation of 0.4. Thus, predictions of new drug combinations in neoplasms have considerably higher value and confidence. ATC codes: *G02CC01;C01EB16;M01AE01;M02AA13;R02AX02;N02AJ08;N02AJ19;M01AE51, **B01AC06;C07FX04;C10BX04;M01BA03;N02BA71;C10BX02;B01AC56;N02AJ07;N02AJ02;N02BA01;C10BX05;N02BA51;A01AD05;C10BX01;C07FX03;N02AJ18;C10BX12;C10BX08;C10BX06;C07FX02, ***G02CC01;C01EB16;M01AE01;M02AA13;R02AX02;N02AJ08;N02AJ19;M01AE51, **** C09BX03;C10BX04;C09BB05;C09AA05;C10BX18;C10BX17;C09BB07;C09BX05;C10BX06;C09BA05, *****C09BX03;C10BX04;C09BB05;C09AA05;-C10BX18;C10BX17;C09BB07;C09BX05;C10BX06;C09BA05

| cardiovascular | | | | |
|---|---|---|---|---|
| drugA name | drugB name | prediction probability | drugA ATC codes | drugB ATC codes |
| Top ten (potentially effective) drug combinations | | | | |
| IBUPROFEN | HYALURONIC ACID | 0.97201 ± 0.00838 | * | S01KA51;S01KA01;D03AX05;M09AX01;R01AX09 |
| PRENYLAMINE | HYALURONIC ACID | 0.96814 ± 0.015 | C01DX52;C01DX02 | S01KA51;S01KA01;D03AX05;M09AX01;R01AX09 |
| ALPROSTADIL | HYALURONIC ACID | 0.96577 ± 0.01473 | C01EA01;G04BE01 | S01KA51;S01KA01;D03AX05;M09AX01;R01AX09 |
| UREA | PRENYLAMINE | 0.94927 ± 0.03799 | D02AE01;B05BC02;D02AE51 | C01DX52;C01DX02 |
| ALPROSTADIL | RANIBIZUMAB | 0.93617 ± 0.02816 | C01EA01;G04BE01 | S01LA04 |
| PROBUCOL | HYALURONIC ACID | 0.93319 ± 0.02795 | C10AX02 | S01KA51;S01KA01;D03AX05;M09AX01;R01AX09 |
| ACETYLSALICYLIC ACID | PALIFERMIN | 0.92995 ± 0.04482 | ** | V03AF08 |
| ALPROSTADIL | TRANEXAMIC ACID | 0.90504 ± 0.07249 | C01EA01;G04BE01 | B02AA02 |
| GERANYL DIPHOSPHATE | PITAVASTATIN | 0.90362 ± 0.14173 | NaN | C10AA08 |
| IBUPROFEN | PALIFERMIN | 0.90244 ± 0.12429 | *** | V03AF08 |
| Bottom ten (potentially adverse) drug combinations | | | | |
| ENALAPRIL | CILAZAPRIL | 0.0 ± 0.0 | C09BA02;C09BB06;C09BB02;C09AA02 | C09BA08;C09AA08 |
| RAMIPRIL | FOSINOPRIL | 0.0 ± 0.0 | **** | C09BA09;C09AA09 |
| RAMIPRIL | QUINAPRIL | 0.0 ± 0.0 | ***** | C09AA06;C09BA06 |
| QUINAPRIL | CILAZAPRIL | 0.0 ± 0.0 | C09AA06;C09BA06 | C09BA08;C09AA08 |
| FOSINOPRIL | BENAZEPRIL | 0.0 ± 0.0 | C09BA09;C09AA09 | C09BB13;C09BA07;C09AA07 |
| DOFETILIDE | DOBUTAMINE | 0.0 ± 0.0 | C01BD04 | C01CA07 |
| PIRBUTEROL | PROCAINAMIDE | 0.0 ± 0.0 | R03CC07;R03AC08 | C01BA02 |
| ISOETHARINE | PROCAINAMIDE | 0.0 ± 0.0 | R03CC06;R03AC07 | C01BA02 |
| FLUCYTOSINE | DOFETILIDE | 0.0 ± 0.0 | J02AX01;D01AE21 | C01BD04 |
| DOFETILIDE | TUBOCURARINE | 0.0 ± 0.0 | C01BD04 | M03AA02 |

**Table S13:** Top ten (potentially effective) and bottom ten (potentially adverse) predicted new drug combinations for nervous system diseases. Predictions are shown for drug combinations with at least one drug with an ATC code starting with N for nervous system diseases. The mean and standard deviation of prediction probabilities over the five-fold cross-validation models are displayed. Top potentially effective predictions in nervous system diseases for anti-nervous system diseases drugs score between 0.65 and 0.78 with a relatively high standard deviation of 0.4. Thus, predictions of new drug combinations in neoplasms have considerably higher value and confidence. ATC codes: *S01KA51;S01KA01;D03AX05;M09AX01;R01AX09, **B01AC06;C07FX04;C10BX04;M01BA03;N02BA71;C10BX02;B01AC56;N02AJ07;N02AJ02;N02BA01;C10BX05;N02BA51;A01AD05;C10BX01;C07FX03;N02AJ18;C10BX12;C10BX08;C10BX06;C07FX02

| nervous system | | | | |
|---|---|---|---|---|
| drugA name | drugB name | prediction probability | drugA ATC codes | drugB ATC codes |
| Top ten (potentially effective) drug combinations | | | | |
| METIXENE | PILOCARPINE | $0.95558 \pm 0.02009$ | N04AA03 | N07AX01;S01EB01;S01EB51 |
| TRIHEXYPHENIDYL | PILOCARPINE | $0.95558 \pm 0.02009$ | N04AA01 | N07AX01;S01EB01;S01EB51 |
| CRYPTENAMINE | PILOCARPINE | $0.95558 \pm 0.02009$ | NaN | N07AX01;S01EB01;S01EB51 |
| ATROPINE | PILOCARPINE | $0.95558 \pm 0.02009$ | S01FA01;A03CB03;A03BA01 | N07AX01;S01EB01;S01EB51 |
| HOMATROPINE METHYLBROMIDE | PILOCARPINE | $0.95558 \pm 0.02009$ | A03CB04;A03BB06 | N07AX01;S01EB01;S01EB51 |
| CAFFEINE | GAMMA-AMINOBUTYRIC ACID | $0.95373 \pm 0.042$ | D11AX26;R03DA20;V04CG30;N06BC01 | NaN |
| SALICYLIC ACID | HYALURONIC ACID | $0.92829 \pm 0.07858$ | D01AE12;N02BA04;N02BA12;S01BC08 | * |
| SALICYLIC ACID | ALEGLITAZAR | $0.92648 \pm 0.05677$ | D01AE12;N02BA04;N02BA12;S01BC08 | NaN |
| SALICYLIC ACID | BEZAFIBRATE | $0.92272 \pm 0.05338$ | D01AE12;N02BA04;N02BA12;S01BC08 | C10AB02 |
| ACETYLSALICYLIC ACID | BEZAFIBRATE | $0.92272 \pm 0.05338$ | ** | C10AB02 |
| Bottom ten (potentially adverse) drug combinations | | | | |
| NITRAZEPAM | ELLAGIC ACID | $0.0002 \pm 0.00027$ | N05CD02 | NaN |
| SORAFENIB | TOFISOPAM | $0.0002 \pm 0.00026$ | L01EX02 | N05BA23 |
| MEPERIDINE | SPARFOSIC ACID | $0.00019 \pm 0.00026$ | N02AB02;N02AB72;N02AB52;N02AG03 | NaN |
| VANDETANIB | BROTIZOLAM | $0.00019 \pm 0.00029$ | L01EX04 | N05CD09 |
| SUNITINIB | RAMELTEON | $0.00018 \pm 0.00023$ | L01EX01 | N05CH02 |
| IMIPRAMINE | ALVOCIDIB | $0.00018 \pm 0.00025$ | N06AA02 | NaN |
| MEPERIDINE | OUABAIN | $0.00015 \pm 0.00021$ | N02AB02;N02AB72;N02AB52;N02AG03 | C01AC01 |
| MEPERIDINE | ACETYLDIGITOXIN | $0.00015 \pm 0.00021$ | N02AB02;N02AB72;N02AB52;N02AG03 | C01AA01 |
| MEPERIDINE | BRETYLIUM | $0.00015 \pm 0.00021$ | N02AB02;N02AB72;N02AB52;N02AG03 | C01BD02 |
| MEPERIDINE | DESLANOSIDE | $0.00015 \pm 0.00021$ | N02AB02;N02AB72;N02AB52;N02AG03 | C01AA07 |
